# ^1^H NMR chemical exchange techniques reveal local and global effects of oxidized cytosine derivatives

**DOI:** 10.1101/2021.12.14.472563

**Authors:** Romeo C. A. Dubini, Eva Korytiaková, Thea Schinkel, Pia Heinrichs, Thomas Carell, Petra Rovó

**Author notes:** Contributed equally to this work (Thea Schinkel, Pia Heinrichs).

## Abstract

5-carboxycytosine (5caC) is a rare epigenetic modification found in nucleic acids of all domains of life. Despite its sparse genomic abundance, 5caC is presumed to play essential regulatory roles in transcription, maintenance and base-excision processes in DNA. In this work, we utilize nuclear magnetic resonance (NMR) spectroscopy to address the effects of 5caC incorporation into canonical DNA strands at multiple pH and temperature conditions. Our results demonstrate that 5caC has a pH-dependent global destabilizing and a base-pair mobility enhancing local impact on dsDNA, albeit without any detectable influence on the ground-state B-DNA structure. Measurement of hybridization thermodynamics and kinetics of 5caC-bearing DNA duplexes highlighted how acidic environment (pH 5.8 and 4.7) destabilizes the double-stranded structure by ~10-20 kJ mol^−1^ at 37 °C when compared to the same sample at neutral pH. Protonation of 5caC results in a lower activation energy for the dissociation process and a higher barrier for annealing. Studies on conformational exchange on the *μ*s time scale regime revealed a sharply localized base-pair motion involving exclusively the modified site and its immediate surroundings. By direct comparison with canonical and 5-formylcytosine (5fC)-edited strands, we were able to address the impact of the two most oxidized naturally occurring cytosine derivatives in the genome. These insights on 5caC’s subtle sensitivity to acidic pH contribute to the long standing questions of its capacity as a substrate in base excision repair processes and its purpose as an independent, stable epigenetic mark.

## Introduction

Apart from the four canonical bases, DNA may also contain modified versions of cytosine, thymine or adenosine nucleosides.^1,2^ Such naturally occurring DNA modifications are broadly named epigenetically modified bases, and they constitute an additional regulatory layer by extending genomic complexity, and have been shown to play several crucial roles.^3^ The discovery of ten-eleven translocation (TET)-induced oxidation of 5-methylcytosine (5mC) to 5-hydroxymethylcytosine (5hmC) has been a key catalyst for the exploration and subsequent characterization of both 5-formylcytosine (5fC) and 5-carboxycytosine (5caC).^4^

5fC and 5caC retain a yet undefined number of functional roles, besides being intermediates within the active demethylation pathway,^5,6^ as they represent genomically stable, semi-permanent modifications with clearly defined tissue distributional patterns.^7–9^ Both have been reported as abnormally abundant in prostate, breast and plasma cells cancer.^10–13^ Their biological significance is not limited to the initial appearance and progression of diseases, since 5fC and 5caC are transiently accumulated during lineage specification of neural stem cells (NSCs) in culture and *in vivo*, ^14^ reduce the rate and substrate specificity of RNA polymerase II transcription,^15^ or can be selectively recognized by specialized proteins.^16^

Despite extensive research efforts in recent years, it is yet unclear how reader proteins recognize 5fC and 5caC with high specificity and selectivity regardless of their sparse genomic abundance and their chemical similarity to canonical, methylated or hydroxymethylated cytosines. 5fC has been more extensively studied compared to its carboxylated counterpart. Recent research endeavors clarified initial contradictory reports about its impact on structure and stability, leaning towards the notion that 5fC does not affect the B-DNA form.^17–19^ As a result of the destabilizing effect that formylation imparts on the C–G base pair, 5fC was found to facilitate melting and hinders annealing, although without affecting the structure to any measurable extent, an aspect which has been investigated via FT-IR and NMR spectroscopy. ^20,21^

From a structural perspective, it has been recently suggested that 5caC is able to induce a kink in dsDNA, and such geometric alteration has been deemed essential for recognition and enzymatic action by Thymine DNA Glycosylase (TDG).^22,23^ On the other hand, disagreement arises concerning its impact on dsDNA melting and annealing, with studies reporting inconsistent de-/stabilization-related properties even when studied under nearly identical conditions. ^24–26^ While 5fC-induced weakening of the base pair was found to be independent of the mildly basic or acidic conditions naturally occurring within different tissues and/or organisms, with the introduction of the titrable carboxyl group, a pH-dependent behavior emerges for 5caC.^23,26^ Protonation/deprotonation events of the carboxyl group induce 5caC to act as an electron-withdrawing group (EWG)/electron-donating group (EDG), respectively. These aspects have been reported to be biologically significant. Indeed, the carboxyl group protonation state has a subtle impact on hydrogen bonding, a behavior that was rationalized via the pKa values of the two solvent-exposed sites: the nitrogen atom N3 and the carboxyl group itself.^21,23,27^ Correspondingly, activity studies on DNA polymerases also concluded that 5caC acts as a base-pair mismatch during DNA replication, signaling that protonated 5caC is a highly destabilizing entity in the context of dsDNA.^28^ In addition, structural biology investigations have suggested that the degree of protonation of 5caC’s carboxyl group might be a key factor in the mechanism of excision operated by TDG.^23^ Even though other studies have considerably advanced our understanding of epigenetically modified DNA bases within the context of nucleic acids, a number of aspects remain unsettled. In this paper, we seek to shed light on the extent to which 5caC–edited dsDNA’s structural features and kinetics-related phenomena deviate from its formylated and canonical equivalents. By employing state-of-the-art methodologies and analytical frameworks in solution-state NMR spectroscopy, we aimed at providing a non-invasive, label-free and site-specific description of the structure and dynamics of these samples using three distinct approaches. Firstly, we compared the impact of the 5caC modification on dsDNA at pH 7.0, 5.8 and 4.7 with respect to 5fC and canonical C by measuring ^1^H, ^13^C and ^15^N chemical shift perturbations. Secondly, we applied temperature-dependent ^1^H chemical exchange saturation transfer (CEST) experiments to assess the impact of 5caC on dsDNA melting and annealing processes providing site-specific parameters for dissociation 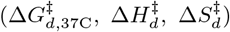 and association kinetics 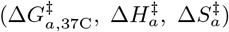 and for melting thermodynamics 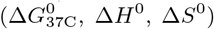.^20,29^ Thirdly, we explored the *μ*s timescale exchange kinetics via ^1^H *R*_1*ρ*_ relaxation dispersion (RD), specifically targeting potential base-specific motions. Such dynamic processes are expected to be absent in undamaged, non-mismatched DNA helices which are not interacting with a reader protein or enzyme.^30,31^

Collectively, our findings reveal that 5caC’s pH-induced chameleonic behavior is not due to any permanent structural changes, opposite to previously reported results. ^22,23^ In fact, we observed that the repercussions of 5caC and 5fC incorporation into otherwise canonical dsDNA are uniquely perceptible in the context of conformational dynamics affecting at least two distinct timescales: a slower one (tens of milliseconds) and a faster one (hundreds of microseconds). In addition, we found that carboxycytosine does noticeably affect dsDNA annealing and melting phenomena when exposed to progressively lower pH conditions. Our unified analysis provides a highly comparable and comprehensive overview of the mechanistic details involving both global (such as energetics of DNA strands association and dissociation) and local (site specific motions involving a single base-pair) dynamics phenomena, and sets the stage for future studies of protein-DNA interactions.

## Results

We considered the pH-dependent structural features, thermodynamic stability, and site-specific dynamics of a self-complementary 12mer homo-carboxylated DNA with the sequence of 5’-GCGATXGATCGC-3’ where X stands for 5caC (Figure 1). The corresponding samples were named caC_7.0_, caC_5.8_, and caC_4.7_ indicating the pH at which they were studied. In order to evaluate the influence of cytosine carboxylation in a broader context, we compare the chemical shifts and the NMR-derived thermodynamics, kinetics, and dynamics parameters to analogous values obtained for canonical (C_7.0_) and 5fC-modified (fC_7.0_) samples featuring a sequence identical as what we considered here, which were previously reported in Ref. 20.

**Figure 1.**
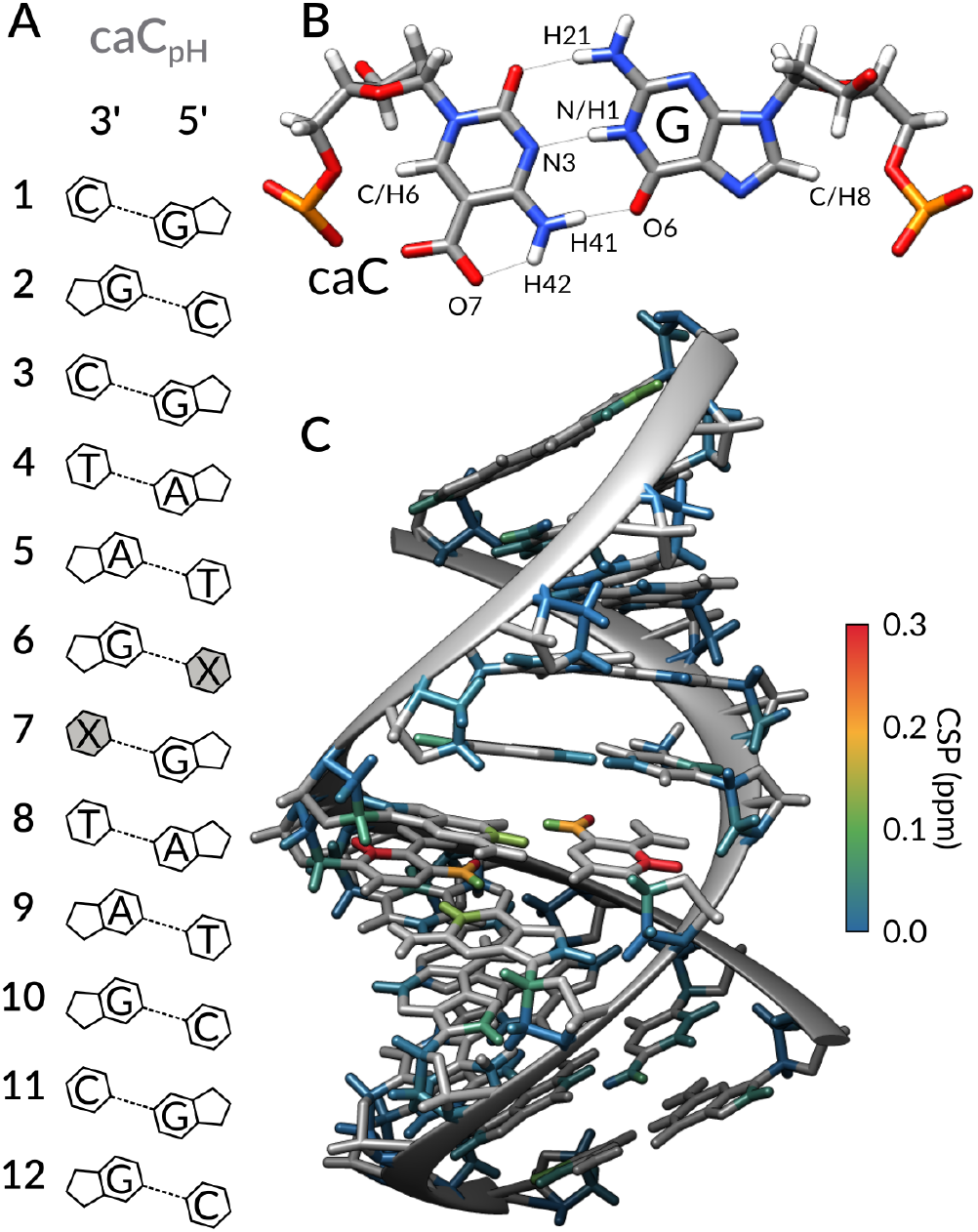
A) DNA sequence of caC_pH_’s model sequence. The modified cytosine nucleoside is highlighted in gray, marked as X. (B) Structural model showing the expected 5caC–G basepair conformation, consistent with ^1^H, ^13^C and ^15^N chemical shift values. (C) Absolute CSP values comparing samples at pH 7.0 and 4.7 are schematically displayed onto the B-DNA structural model of caC_pH_.

### Structural impact by chemical shifts analysis

To spot any unusual features induced by inclusion of carboxylated cytosine within otherwise canonical DNA, we recorded a comprehensive set of homo-(^1^H-^1^H NOESY) and heteronuclear (^1^H-^13^C HSQC, ^1^H-^15^N SOFAST HMQC) 2D spectra for resonance assignment and chemical shift analysis purposes. ^32^

In contrast to crystallographic results reported in ref. 22 and 23, our results indicate that 5caC nucleobase does not induce any detectable, permanent deviation from the canonical B-DNA structure. At pH 7.0, when compared to the 5fC–G base-pairing interactions, 5caC–G appears to be equally well-tolerated within a canonical double-helix architecture: all of ^1^H-^13^C and ^1^H-^15^N cross-peaks are superimposable among the canonical, and 5fC and 5caC-containing 12mer DNA constructs, with the sole exception represented by those nuclei in direct proximity to the epigenetic modification (Fig. 2, S1, S2).

**Figure 2.**
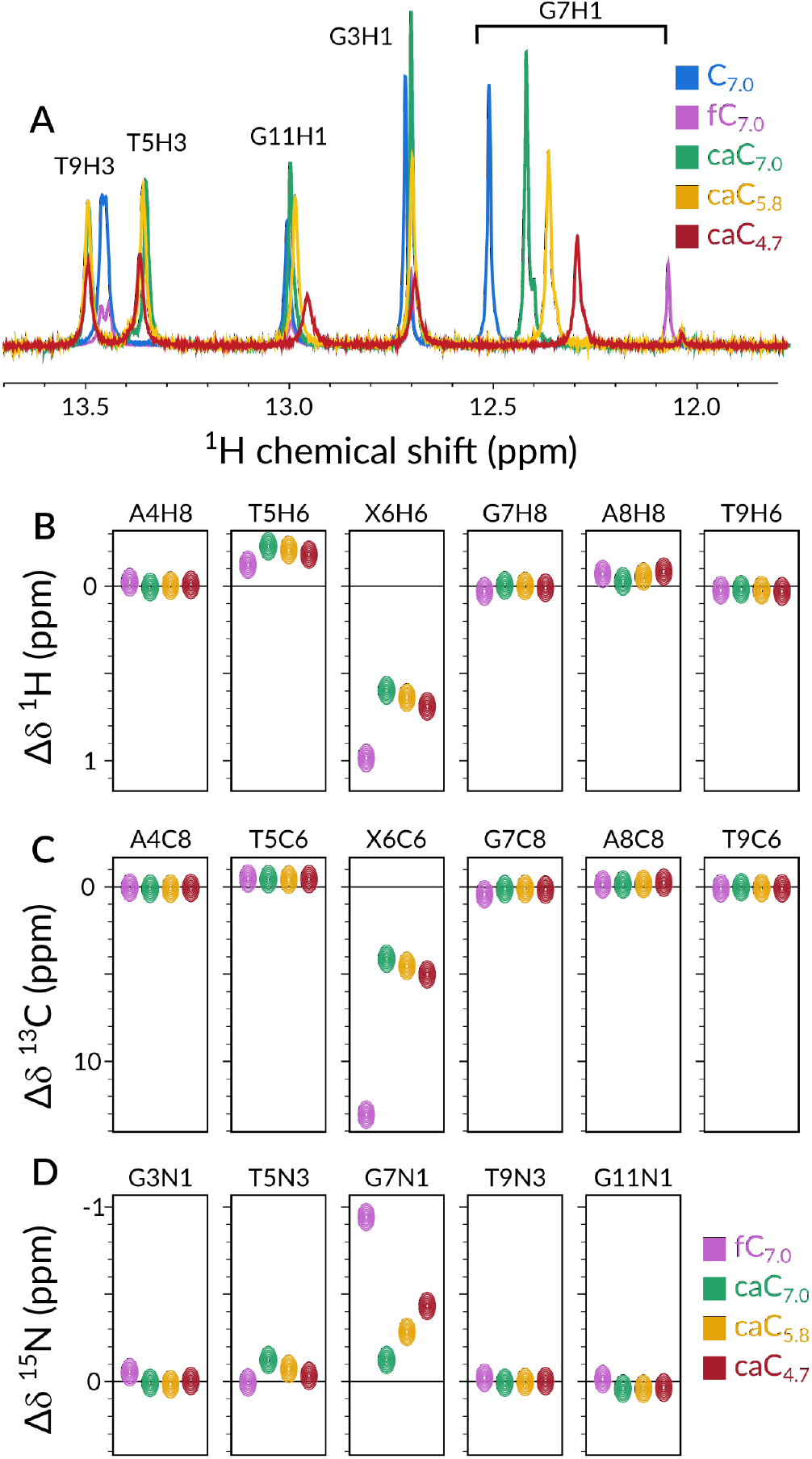
Comparison of chemical shift perturbations Δ*δ* for representative imino ^1^H (A), aromatic ^1^H (B), aromatic ^13^C (C), and imino ^15^N (D) resonances with respect to the chemical shifts of the canonical 12mer sample (blue spectrum in A, black horizontal line in B–D. Magenta, green, yellow and red symbols represent fC_7.0_, caC_7.0_, caC_5.8_ and caC_4.7_, respectively. A full comparison of all comparable chemical shifts between the five samples is displayed in Fig. S7–S10.

An analogous comparison of caC_7.0_ with caC_5.8_ and caC_4.7_ points to no detectable structural change upon acidification. ^1^H, ^13^C, and ^15^N resonances, ^3^J_HH_ couplings and ^1^H-^1^H NOESY cross-peak patterns suggest that carboxylated cytosine leaves the B-DNA structure entirely unperturbed at both neutral and acidic pH values, resulting in no substantial structural difference from the canonical or formylated cytosine-bearing constructs in any studied condition (Fig. S3-S6).

Fig. 2 compares the perturbations of representative ^1^H, ^13^C and ^15^N chemical shifts of fC_7.0_, caC_7.0_, caC_5.8_ and caC_4.7_ with respect to the shifts of the canonical 12mer sequence. Fig. 2A shows imino protons, which are sensitive reporters of the strength of base-pairing via hydrogen bonding. Interestingly, the only responsive site to pH changes is G7H1, which reports on the X:G base pair, where X is either C, 5fC or 5caC at the three distinct pH conditions we selected. Substantial chemical shift perturbations are also visible for aromatic ^1^H sites T5H6 and X6H6, ^13^C X6C6 and ^15^N G7N1 (Fig. 2B–D, S7–10). Among those nuclei, the trends suggest that caC_4.7_’s resonances resemble fC_7.0_ the most, while in neutral conditions caC_7.0_ is comparable with canonical C_7.0_.

These pH-induced shifts were most prominent for H-bond forming protons, namely for caC6H41, H42, and G7H1. The nature of the effect is related to the reshuffling of electron densities around the base-pairing atoms due to altered Pauli repulsion between occupied atomic orbitals. ^33^ An increased ^1^H chemical shift is associated with a decreased electron density (^1^H shielding) around the proton and hence with a stronger H-bond, or shorter H–X distance. As the carboxyl group of 5caC becomes progressively protonated with decreasing pH, the intra-base H-bond between caC6H42 and the carboxyl oxygen O7 gets weaker (Δ*δ* = −0.3 ppm), while the interbase H-bond formed by caC6H41 and G7O6 (Fig. 1B) gets stronger (Δ*δ* = +0.1 ppm). Meanwhile, hydrogen bonding between G7H1 and caC6N3 weakens (Δ*δ* = −0.15 ppm) due to the decreased electron density at N3 caused by the EWG properties of the protonated carboxyl group, compensating for the increased base-pairing stability gained by the H41–O6 interaction (Fig. S7–S10). The fine balance between the strengths of the intra- and inter-bases hydrogen bonds in the 5caC–G base pair leads to close to optimal base-pairing both at neutral and mildly acidic conditions, leaving the B-DNA structure unperturbed through the entire studied pH range.

### 5caC’s influence on DNA melting and annealing

A thorough characterization of nucleic acids’ global structural rearrangements, such as folding, melting, annealing and binding should entail a comprehensive analysis of kinetic events occurring on the ms–s time scale. ^34^ CEST experiments have found adoption in modern biomolecular NMR, allowing for quantitative and site-specific determination of population, chemical shift and exchange kinetics of sparsely populated conformations. ^29,35^ When measured in a temperature-dependent fashion, the shift in exchange parameters can reveal atomistic details about the melting thermodynamics and kinetics of the studied system providing unprecedented insights into the molecular processes. In pursuance of the study of 5caC-induced DNA destabilization, we proceeded by recording CEST profiles for the aromatic protons in all caC_pH_ samples in the 55–60 °C range. As an example, Figure 3A displays the melting CEST profiles for C10H6 at three pH values (profiles for other comparable protons can be found in Fig. S11–S18). The appearance of a distinctive secondary dip at increasingly higher temperatures indicates the presence of an alternative conformer, which we identify as the single-stranded conformation (ss-DNA) as per comparative chemical shift analysis.

**Figure 3.**
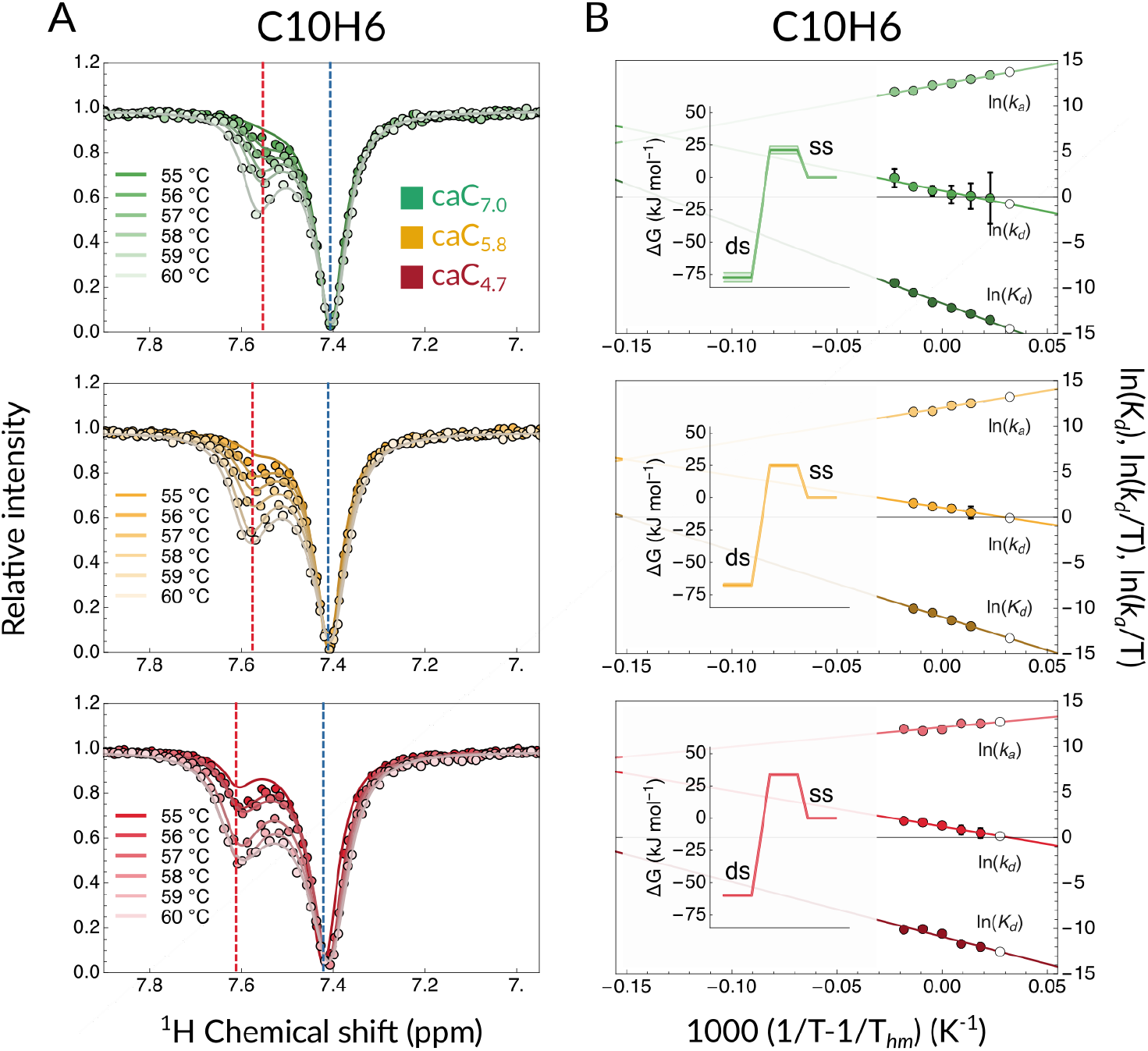
(A) Temperature-dependent CEST melting profiles for proton C10H6. caC_7.0_, caC_5.8_ and caC_4.7_ are shown in shades of green, yellow and red, respectively. Dashed blue and red lines indicate GS and ES chemical shift values. (B) van’t Hoff plots relative to the CEST melting profiles. Shades of green, yellow and red indicate data entries and linear fits for caC_7.0_, caC_5.8_ and caC_4.7_, respectively. White data points represent back-calculated values. Insets present the relevant Gibbs free energy plots at each pH condition at 37 °C.

To obtain a more quantitative comparison, we fit the CEST profiles to a two-state exchange model (dsDNA ⇌ 2 ssDNA) that yielded a numerical estimation of populations (*p*_D_ for dsDNA, 1 – *p*_D_ for ssDNA), exchange kinetics (*k_ex_*), and chemical shifts of the exchanging states at each of the highest temperatures, while we used the lowest temperature of the ensemble for each sample to compare the predicted back–calculated value from the fits to the experimental value. ^1^H longitudinal (*R*_1_) and transverse (*R*_2_) relaxation rates were measured separately at multiple temperatures and used as inputs for the CEST fits assuming that the rate constants of the dsDNA and ssDNA states are the same.

As an example, Figure 3B shows the logarithm of the obtained kinetic rates and equilibrium constants for C10H6. Consistently with results obtained for fC_7.0_ and C_7.0_, two observations can be made: (i) a linear fit could be identified for kinetic rates and equilibrium constants against *T*^-1^, suggesting a single transition state is plausible for the melting and annealing phenomena, and (ii) all sites show thermally activated kinetics with a positive dissociation barrier (Arrhenius behavior) and a negative association barrier (anti-Arrhenius).

In addition, CEST measurements run at multiple distinct temperatures allow for the extraction of Gibbs free energy and related parameters by leveraging the temperature-dependent nature of kinetics and thermodynamic phenomena. In order to achieve an estimation of the degree of protonation-induced destabilization to dsDNA, we applied our recently described methodological framework for the decomposition of traditional CEST output into enthalpic and entropic stability and activation changes.^20^ Succinctly, we assumed the observed dynamic equilibrium is the reversible melting/annealing process of a dsDNA strand into two single-stranded DNA sequences. If the concentration of DNA is known, then kinetics of association (*k_a_*) and dissociation (*k_d_*) can be extracted. From the temperature dependence of *k_a_* and *k_d_*, activation barriers for both process (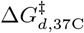 and 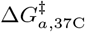) can be derived. Analogously, the ratio between *k_a_*(*T*) and *k_d_*(*T*) rates allows for the determination of the equilibrium dissociation constant, and consequently the quantification of thermodynamic parameters such as 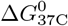.

This analysis allows us to directly compare caC_7.0_, caC_5.8_ and caC_4.7_ not only between them, but also vs. otherwise identical canonical and formylated samples (C_7.0_ and fC_7.0_, respectively), which we previously discussed in ref. 20. In Table 1, we compare three proton reporters across all five samples (a comprehensive table including all proton reportes can be found in Table S1). From a thermodynamic perspective, among the C-modified sample caC_7.0_ scores as the most stable one, which is highly comparable to C_7.0_, as observed by UV/Vis spectroscopy (Fig. S19) as well as in other studies. ^24,27^ Acidification of the buffer to pH 5.8 destabilizes the double-stranded conformer by ~8-10 kJ mol^-1^, and further by another ~2-5 kJ mol^-1^ when the pH is decreased to 4.7. Interestingly, 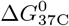 for caC_5.8_’s proton reporters are very similar to those we obtained for fC_7.0_. Kinetics of dissociation data, as expected, support the notion that caC_7.0_ and C_7.0_ require the most energy for undergoing a dsDNA → 2 ssDNA conformational transition. According to this metric, fC_7.0_ is slower in undergoing the melting process when compared to caC_5.8_ and caC_4.7_, as 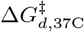 is higher by ~3-6 kJ mol^-1^. Lastly, kinetics of association suggest that caC_4.7_ is the slowest in performing an annealing process, followed by fC_7.0_. Compellingly, 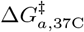 data indicate that the rate of association of caC_7.0_ is ~5-8 kJ mol^-1^ less energetically demanding when compared to C_7.0_, a behavior that can be rationalized considering the EDG nature of the carboxylate substituent (5caC at neutral pH) when compared to a proton (canonical C).

**Table 1.** Thermodynamic and kinetic parameters of the dsDNA melting process obtained from the van’t Hoff and Eyring analysis of the CEST-derived exchange parameters. Errors are given as one standard deviation.

| Sample | | $\Delta G_{37C}^0$<br>kJ mol<sup>-1</sup> | $\Delta G_{d,37C}^\ddagger$<br>kJ mol<sup>-1</sup> | $\Delta G_{a,37C}^\ddagger$<br>kJ mol<sup>-1</sup> |
| --- | --- | --- | --- | --- |
| C<sub>7.0</sub> | C2H6 | 71.1 ± 1.7 | 99.2 ± 2.2 | 28.1 ± 2.1 |
| | T9H6 | 70.2 ± 1.8 | 102.2 ± 1.5 | 31.9 ± 1.5 |
| | C10H6 | 71.3 ± 1.4 | 100.3 ± 1.3 | 29.0 ± 1.3 |
| caC<sub>7.0</sub> | C2H6 | 75.4 ± 7.0 | 98.7 ± 7.8 | 23.3 ± 8.2 |
| | T9H6 | 75.7 ± 0.8 | 99.2 ± 0.9 | 23.4 ± 0.9 |
| | C10H6 | 77.3 ± 3.6 | 98.6 ± 3.2 | 21.3 ± 2.9 |
| caC<sub>5.8</sub> | C2H6 | 66.5 ± 1.5 | 94.4 ± 1.8 | 27.9 ± 1.7 |
| | T9H6 | 65.2 ± 1.2 | 94.9 ± 1.1 | 29.7 ± 1.2 |
| | C10H6 | 67.4 ± 1.4 | 92.4 ± 1.2 | 24.9 ± 1.1 |
| caC<sub>4.7</sub> | C2H6 | 58.6 ± 0.7 | 93.1 ± 0.7 | 34.5 ± 0.9 |
| | T9H6 | 62.1 ± 0.4 | 95.2 ± 0.4 | 33.1 ± 0.5 |
| | C10H6 | 56.8 ± 0.6 | 92.9 ± 0.6 | 36.1 ± 0.6 |
| fC<sub>7.0</sub> | C2H6 | 74.7 ± 10.4 | 99.7 ± 5.6 | 25. ± 5.5 |
| | T9H6 | 64.6 ± 1.2 | 97.1 ± 1. | 32.5 ± 1. |
| | C10H6 | 66.4 ± 1.2 | 97.5 ± 1. | 31.1 ± 1.1 |

In Figure 4 we show the correlation between the dissociation and equilibrium free energy changes across all samples and conditions, for every proton reporter. Here, 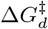 is plotted as a function of Δ*G*^0^ for all five samples across all eligible proton reporters. Data points accounting for caC_7.0_ and C_7.0_ tend to cluster at the upper right-hand corner of the plot. Conversely, 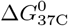 and Δ*G*^‡^*d*, 37C values are substantially decreased whenever fC_7.0_, caC_5.8_ or caC_4.7_ are considered, as elaborated above.

**Figure 4.**
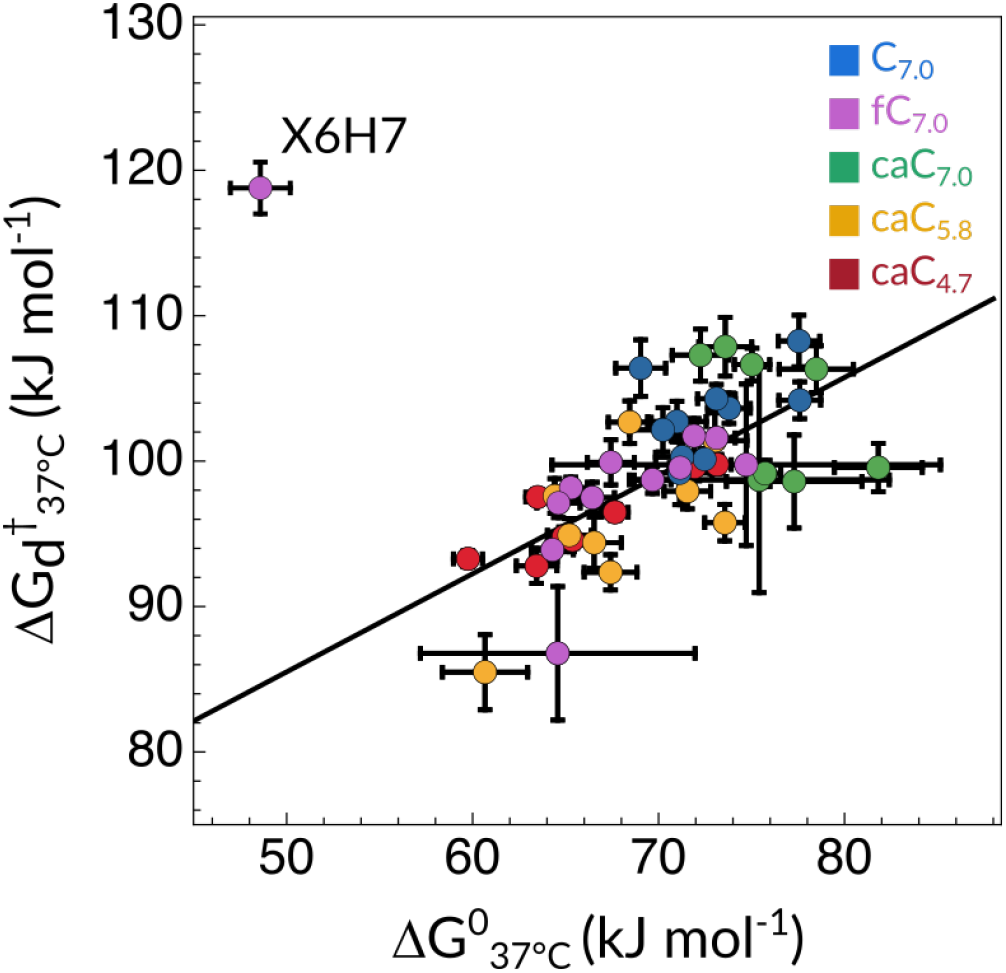
Correlation plot between the changes of the Gibbs free energy of activation for the dissociation process and the equilibrium free energy of previously reported values for fC_7.0_ and C_7.0_ (magenta and blue, respectively) together with 5caC-containing samples (caC_7.0_ as red, caC_5.8_ as yellow and caC_4.7_ as red) at 37 C. fC_7.0_ X6H7 (featuring much higher activation free energies and lower equilibrium energies than the rest of the molecule) is an outlier due to its stable intramolecular hydrogen-bond between the formyl O7 atom and the adjacent amino H42 proton.

### Protonation-induced *μ*s dynamics

^1^H and ^15^N chemical shift values for G7H1/N1 nuclei, both reporters of the centrally positioned 5caC6-G7 base pair’s stability, have evidenced that the protonation state of the exocyclic carboxylic group 5caC has a selective impact on this imino proton resonance (Figure 2A, D), which have long been regarded as key indicators of hydrogen bond strength.^24,36^ Because of this observation, we aimed at investigating whether this protonation-induced, localized weakening is accompanied by increased probability of local fast time-scale motions. Despite the rich ensemble of motions occurring across the nanosecond and the milliseconds intervals, both experimental data and molecular dynamics simulation studies have shown that, provided no mismatch or abasic site is present, intermediate (microseconds) time scale dynamics are normally absent in B-DNA helices. ^30^

Our CEST-based kinetic and thermodynamic analysis has established that caC_5.8_, caC_4.7_ and fC_7.0_ destabilize the double-stranded DNA structure without any apparent static, persistent impact on its helical architecture. In order to investigate the presence of a potentially localized conformational exchange which might contribute to the destabilization, we interrogated the faster, microsecond time scale by applying ^1^H R_1*ρ*_ relaxation dispersion (RD) methods. ^34,37^ We measured *R*_1*ρ*_ RD profiles in the 1 to 10 kHz range at 55 °C, for each sample in identical conditions.

In Figure 5 we show X6H6 (where X = C, 5fC or 5caC, depending on the sample under current consideration) ^1^H *R*_1*ρ*_ RD profiles measured at 55 °C, a condition that ensures that the melting process is still rather sparse and infrequent. The data set recorded for caC_7.0_, caC_5.8_ and C_7.0_ are best fit to a no-exchange model, resulting in flat profiles (black lines). Conversely, profiles for caC_4.7_ and fC_7.0_ fit best to the exchange model, exposing a chemical exchange contribution to its *R*_2_ relaxation rate. Such motions, consistent with a *τ_ex_* in the order of hundreds of microseconds, fall within the intermediate exchange regime and are approximately two orders of magnitude faster compared to the overall melting process as characterized by our CEST measurements. This result is especially interesting when comparing the H6 proton of the modified base in samples caC_4.7_ and fC_7.0_ to any other available ^1^H nucleus. Apart from A8H8, which appears to also engage in a *μ*s time scale motion (likely in response to 5caC’s motion) no other profile is consistent with an exchange phenomenon on this interval (Fig. S20-S24). In other words, this motions appears to be sharply localized, leaving all other bases unaffected.

**Figure 5.**
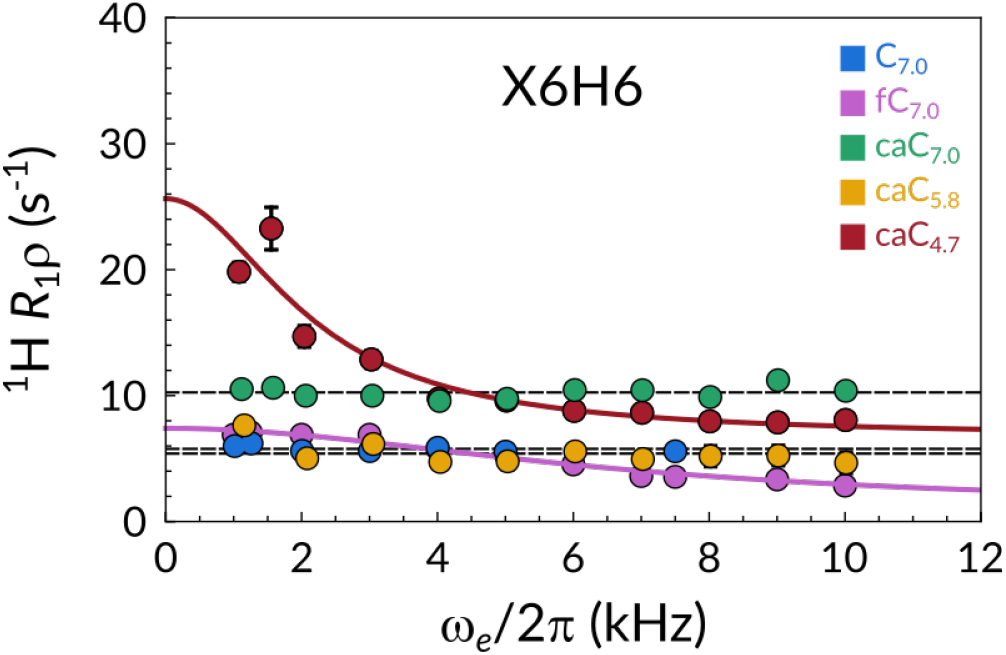
R_1*ρ*_ relaxation dispersion profiles of C_7.0_ (blue), fC_7.0_ (magenta), caC_7.0_ (green), caC_5.8_ (yellow) and caC_4.7_ (red) recorded at 55 °C. Lines represent best fits either to models accounting for (color-coded) or discounting chemical exchange (black).

This result confirms our previous speculations on the mechanism by which 5fC (and by extension protonated 5caC) weakens the dsDNA conformer. ^20^ Deprotonated (or not protonated to a sufficient extent) 5caC proved itself to be either a neutral or even a stabilizing factor in terms of base-pair strength and agrees well both with NOESY-derived chemical shift values and the literature. Instead, whenever the pH of the buffer is sufficiently acidified, the weakening of the 5caC-G hydrogen bond induces a chemical exchange process in the intermediate microsecond time scale. The fact that this motion is sharply localized at the modified site seems to suggest that the protonation of the exocyclic carboxyl moiety and the consequent lowered basicity of the hydrogen-bonded N3 atom could be deemed responsible for generating a sparse and transitory fraying event in the middle of the DNA strand, presumably synergistic with the global destabilization effect on the whole structure.

## Discussion

In pursuance of understanding the role of 5caC, we addressed the structural and dynamic features of oligomeric DNA double strands carrying a single version of such modification on each strand. Incorporation of carboxycytosine into DNA is a naturally occurring phenomenon that has been mainly discussed within two biological frameworks: its recently devised (and to date mostly obscure) semipermanent epigenetic role, and as a DNA lesion that is undergoes the base excision repair (BER) process.^2-4,38^

### Considerations on 5caC protonation sites

In their report on divergent mechanism of enzymatic excision for 5caC and 5fC, Maiti *et al*. have elaborated how, in a TDG-DNA complex, nitrogen N3 of 5caC (Fig. 1B) is likely more basic than the carboxyl group, and thus undergoes protonation before the carboxylic exocyclic moiety does. ^23^ In contrast, despite assigning a pK_a,COO_- of 4.7 and pK_a,N3_ of 2.1 for the isolated nucleoside, a distinct infrared spectroscopy and quantum mechanical analysis suggests that, when in the context of a dsDNA strand, the first protonation site is the carboxyl group.^26^ Our results support the latter idea. In Fig. 2A, we show that no other base pair is affected by the pH change over the entire 7.0-4.7 interval. Indeed, T9H3, T5H3 and G3H1 display negligible chemical shift differences, while G11H1 shifts upfield by ~0.05 ppm likely due to its proximity to the fraying ends of the oligomeric model system. In sharp contrast, the signal reporting on G7H1 consistently shifts upfield with decreasing pH. While this preference for protonation of the weaker base (i.e., 5caC’s carboxylate group over N3) is apparently counterintuitive, we reason that in a dsDNA setting the N3 site is well protected from the solvent environment, both for steric and electrostatic reasons.

### 5fC and 5caC as semi-permanent modifications

For long, 5fC and 5caC have been mainly considered as transient intermediates within the active demethylation pathway. ^39^ However, research towards their capacity as standalone epigenetic marks has gained increasingly more traction. For instance, both 5fC and 5caC overlap with H3K4me1 marked regions, associated with active transcription. ^40^ Also, several developmental and metabolic related genes show 5fC enrichment on promoters before gene upregulation, ^41^ while 5caC has been reported to transiently accumulate at promoter regions preceding gene expression during lineage specification and differentiation.^42^ Lastly, a number of cancerous diseases are correlated with a significant enrichment of such oxidized cytosine epigenetic modifications.^10–13^

From a structural and conformational perspective, both 5fC and 5caC have been reported to induce structural changes localized in the proximity of the modified nucleoside, ^22,23^ while the pH dependence of 5caC characteristics on dsDNA stability have been previously discussed in the context of short oligomers carrying several clustered modifications.^21,27^ In the context of longer (90 bp) DNA strands, Ngo *et al*. reported a three-fold enhancement of cyclization rate for 5fC containing strands, while the sample carrying 5caC at pH 8.0 showed no detectable difference when compared to canonical cytosine. ^43^

In contrast to Ref. 22,23, our results indicate that, identically to 5fC, 5caC does not induce any permanent structural modification detectable by NMR spectroscopy at any pH condition under consideration (Fig. 2, S2-10). In fact, in analogy to 5fC, 5caC was found to affect dsDNA melting and annealing equilibrium and kinetics, rather than B-DNA average structure. On average, and across all sites, caC_5.8_ and caC_4.7_ resemble fC_7.0_, both thermodynamically and kinetically, when it comes to annealing and melting. In contrast, and consistently with previous cyclization essays and FRET studies, the behavior of deprotonated 5caC is most similar to canonical cytosine.

5mC oxidation derivatives have been shown to accumulate and persist in a relatively stable state in certain biological contexts. For this reason, they have been suggested to carry out additional roles apart from being intermediates in biochemical pathways. In light of the foregoing, we contend that our results could correlate with the relative abundance of 5caC and 5fC in cancerous tissues. How protonation of the exocyclic carboxyl moiety affects the thermodynamics and kinetics of melting and annealing *in vitro* has been discussed in this and previous reports.^20,21,26,27^ Our results corroborate the protonation-induced destabilization of the double strand, and we speculate that a similar effect might take place in cancerous tissues, triggered by said low pH environment. Cell proliferation is notoriously accelerated in cancer, and the acidic microenvironment where such diseases thrive is widely recognized as a phenotypic trait, making the presence of the two most oxidized cytosine epigenetic derivatives more than circumstantial. ^44,45^

### 5fC and 5caC-driven enzymatic recognition

In mammals, active DNA demethylation takes place via an enzymatic tandem of TET and TDG, which together govern the initial stages of the BER pathway. In order to successfully complete the removal of 5mC, the presence of either 5fC or 5caC are ultimately necessary, as they are substrates for TDG, which generates the abasic site. ^23^ Apart from TET and TDG, several additional proteins are able to selectively recognize, bind and exert their respective enzymatic activity upon 5fC and 5caC. ^12,13,46^ The mechanism by which different reader enzymes selectively recognize cytosine’s epigenetic modifications has long been established as a crucial theme in chemical biology. ^6^ Across the proposed enzymatic mechanisms, many rely on a specific residue to initiate the base-extrusion process into the active site. For instance, the “pinch-push-pull” mechanism proposed for TDG leans on Arg275 to promote the breakage of an X:G base pair, where X = T, 5fC, or protonated 5caC.^47,48^ Alternative studies suggest that partially extruded nucleotide conformations, which are sparse but naturally occurring events, might play a role in recognition and base excision. ^49^ This second mechanism of action seems to agree with DNA replication studies, which highlighted how 5caC:G base-pair can behave as a DNA lesion. ^28^

We believe our R_1*ρ*_ RD data provide one more piece of evidence in this complex puzzle. By observing site selective and spontaneous base-flipping of protonated 5caC:G we provide further experimental evidence that 5fC and protonated 5caC can indeed act as DNA lesions. ^30^ We hypothesize that, albeit undetected in our study, such kinetic processes could be present at physiological temperatures and decisively impact enzyme recognition and mode of action.

## Conclusions

In this work, we have considered the impact of 5caC incorporation into a model dsDNA oligomer (caC_pH_) in three pH conditions, namely at pH 7.0, 5.8 and 4.7. We obtained chemical shift, melting/annealing and *μ*s conformational exchange data which we could reliably compare with our recent study focusing on 5fC and canonical cytosine. Assignment and chemical shift studies on comparable ^1^H, ^13^C and ^15^N nuclei have shown that there is no evidence of a permanent structural change: all caC_pH_ samples, together with C_7.0_ and fC_7.0_, are compatible with a standard B-DNA helical arrangement. CEST-derived kinetic and thermodynamic data suggested that the reduced cohesion of the X6:G7 base pair, evidenced by chemical shift studies, affects the extent to which nearby bases are able to cooperatively stabilize one another. caC_4.7_ and fC_7.0_ emerged as the most destabilized samples of the cohort, while caC_7.0_ and C_7.0_ showed remarkably similar properties overall.

caC_4.7_ and fC_7.0_ also revealed a detectable chemical exchange process at or in the proximity of the modified nucleoside X6 on the *μ*s time scale. The data hereby presented indicate that 5caC’s impact on B-DNA is only evident through the lenses of conformational dynamics, as protonation of the exocyclic carboxyl moiety affects the melting-annealing equilibrium as well as induces sparse and localized *μ*s time scale base-pair dynamics. We believe our findings are relevant in the context of several open questions concerning this sparse epigenetic mark. Future investigations may consider studying protein-DNA interactions, featuring isotopic labelled 5fC or 5caC nucleosides in order to unravel the exact mechanistic details of the interaction between TET, TDG (and other enzymes) and cytosine’s oxidized derivatives.

## Materials and methods

### Sample preparation

The cadC-phosphoramidite (cadC-PA) and subsequently the modified dsDNA samples caC_pH_ were prepared via phosphoramidite chemistry as previously reported. ^50^ Solid phase synthesis of oligonucleotides containing cadC was performed on an ABI 394 DNA/RNA synthesizer (Applied Biosystems) using standard DNA synthesis conditions with a cartridge scale of 1 *μ*mol. The phosphoramidites Bz-dA, Ac-dC, iBu-dG and dT as well as the PS carriers were purchased from LinkTechnologies. For the reaction of the cadC-PA a coupling time of 180 s was applied. The terminal DMT protecting group was cleaved after DNA synthesis on the synthesizer. Basic and acidic deprotection of all oligonucleotides was performed according to literature. ^50^ Purification of the oligonucleotides was achieved with a HPLC system (Agilent 1260 Infinity II 400 bar pump and a Agilent 1260 Infinity II VWD detecting at 260 nm) applying a buffer system of 0.1 M triethylammonium acetate in water (buffer A) and 0.1 M triethylammonium acetate in 80 % aqueous MeCN (buffer B), a gradient of 0% - 30% buffer B in 45 min and a flow rate of 5.0 mL/min. As stationary phase Nucleodur columns (250/10 mm, C18ec, 5 *μ*m) from Macherey-Nagel were used. Purified oligonucleotides were analyzed by MALDI-TOF (Bruker Autoflex II). Quantification of oligonucleotides was performed via UV/Vis spectroscopy with a NanoDrop ND-1000 spectrophotometer at 260 nm. Samples caC_7.0_ and caC_5.8_ were dissolved in aqueous buffers consisting of 15 mM Na_2_HPO_4_/NaH_2_PO_4_ (pH 7.0 and 5.8, respectively), 25 mM NaCl in H_2_O. Sample caC_4.7_ was prepared by titrating a 1 M HCl solution into the same buffer described above. The thermal stability of the buffer between room temperature and 60 °C was ascertained by pH-meter measurements. Annealing was performed by heating the dsDNA-containing buffer solution to 90 °C for 5 minutes and slowly cooling it to 5 °C in approx. 90 minutes, after which it was allowed to return to room temperature. Then, the NMR sample was prepared with the addition of 0.02% NaN_3_, 25 *μ*M DSS and 5% D_2_O, resulting in final sample concentrations of ~ 0.66 mM for all samples, as determined via UV spectrophotometric measurements at 260 nm using the extinction coefficient calculated via the nearest neighbor approximation.

### UV/Vis spectroscopy

UV/Vis melting profiles of the oligonucleotides were measured at 260 nm with a JASCO V-650 UV/Vis spectrophotometer between 20 and 85 °C (scanning rate of 1 °C/min), and each sample was measured four times. Samples were placed into 100 *μ*L cuvettes and diluted with the same Na_2_HPO_4_/NaH_2_PO_4_, NaCl aqueous buffer as used in the NMR experiment. Before each measurement, a layer of mineral oil was placed on the surface of the sample in order to prevent water evaporation. caC_7.0_ was measured at four concentrations (1.25, 2.50, 5.00, and 10.00 *μ*M), while fC_7.0_ and C_7.0_ were measured as described in Ref. 20. All concentration values yielded absorption values within the linear range of the spectrometer.

### NMR spectroscopy

All experiments were performed on Bruker Avance III spectrometer operating at a ^1^H Larmor frequency of 800 MHz (corresponding to a magnetic field of 18.8 T) equipped with a 5 mm triple-resonance cryogenically cooled TCI probe. Standard 2D NOESY (mixing time 250 ms) spectra were recorded at 37 °C for resonance assignment. Natural abundance ^1^H-^13^C and ^1^H-^15^N HSQC and HMQC spectra were recorded using standard fast-pulsing pulse sequences. ^51^ Site-selective spin relaxation measurements were performed following the SELOPE scheme; these included ^1^H CEST, on-resonance ^1^H R_1*ρ*_, recorded either with a single spin-lock strength of 10 kHz, or as an entire RD profile ranging from 1 to 10 kHz, and ^1^H R_1_ experiments at temperatures between 37 and 60 °C. The employed R_1*ρ*_ and ^1^H CEST pulse sequences have been modified from Schlagnitweit *et al*..^37^ CEST profiles, pseudo-2D ^1^H R_1_ and pseudo-3D R_1*ρ*_ experiments were performed, processed and analyzed as previously described. ^20^

## Supporting Information Available

**Supplementary Figures**: ^1^H-^1^H, ^1^H-^13^C and ^1^H-^15^N 2D correlation spectra, Schematic comparison of ^1^H, ^13^C and ^15^N chemical shift perturbations, Concentration-dependent UV/Vis melting analysis, Temperature-dependent CEST profiles and van’t Hoff plots for all available sites, R_1*ρ*_ relaxation dispersion profiles for all samples. **Supplementary Tables**: CEST-derived thermodynamic and kinetic parameters for all samples. This material is available free of charge via the Internet at http://pubs.acs.org.

## Acknowledgement

We thank Markus Müller for valued discussions and Felix Xu for assistance in the measurement of UV/Vis melting profiles. This work was supported in part by the Deutsche Forschungsgemeinschaft (DFG, German Research Foundation) – SFB 1309 – 325871075, EU-ITN LightDyNAmics (ID: 765266), the ERC-AG EpiR (ID: 741912), the Center for NanoScience, the Excellence Clusters CIPSM, and the Fonds der Chemischen Industrie. Funding for open access charge: the SFB 1309 framework (https://www.sfb1309.de/).

## TOC Graphic

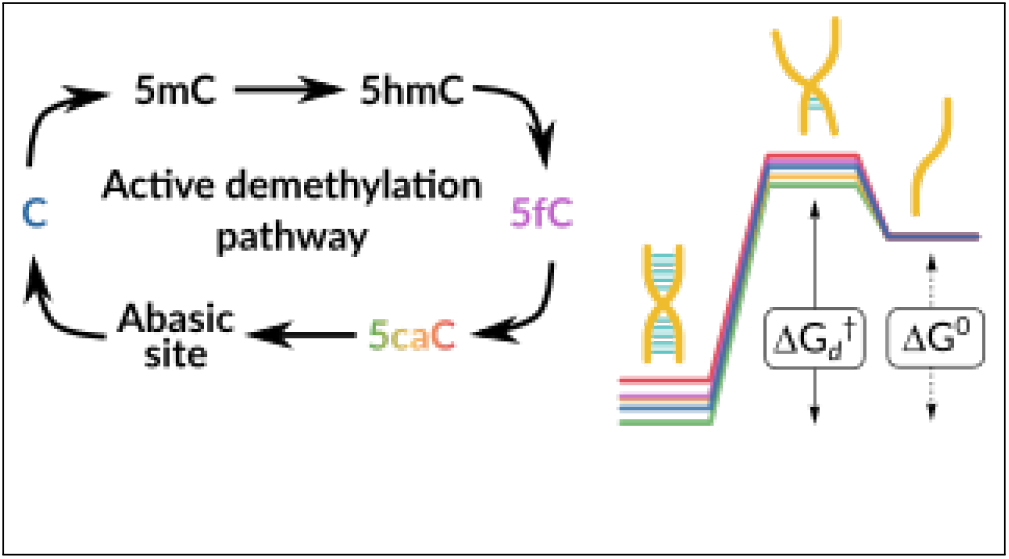

