## Supplementary Material for "^1^H NMR chemical exchange techniques reveal local and global effects of oxidized cytosine derivatives"

§*Contributed equally to this work*

### List of Figures

S1 $^{1}$H-$^{13}$C HSQC aromatic region spectrum of $fC_{7.0}$, $caC_{7.0}$ and $C_{7.0}$ . . . . . . . S3
S2 $^{1}$H-$^{15}$N HMQC imino region spectrum of $fC_{7.0}$, $caC_{7.0}$ and $C_{7.0}$ . . . . . . . . S4
S3 $^{1}$H-$^{15}$N HMQC imino region spectrum of $caC_{7.0}$, $caC_{5.8}$ and $caC_{4.7}$ . . . . . . S5
S4 $^{1}$H-$^{13}$C HSQC aromatic region spectrum of $caC_{7.0}$, $caC_{5.8}$ and $caC_{4.7}$ . . . . . S6
S5 $^{1}$H-$^{1}$H NOESY imino region spectrum of $caC_{7.0}$, $caC_{5.8}$ and $caC_{4.7}$ . . . . . . S7
S6 $^{1}$H-$^{1}$H NOESY aromatic region spectrum of $caC_{7.0}$, $caC_{5.8}$ and $caC_{4.7}$ . . . . S8
S7 Comparison of $^{1}$H chemical shift perturbations for all samples, 1/2 . . . . . . S9
S8 Comparison of $^{1}$H chemical shift perturbations for all samples, 2/2 . . . . . . S10
S9 Comparison of aromatic $^{13}$C chemical shift perturbations for all samples . . . S11
S10 Comparison of aromatic $^{15}$N chemical shift perturbations for all samples . . . S11
S11 Temperature-dependent CEST profiles and van't Hoff plots – C2H6 . . . . . S12
S12 Temperature-dependent CEST profiles and van't Hoff plots – A4H2 . . . . . S13
S13 Temperature-dependent CEST profiles and van't Hoff plots – A4H8 . . . . . S14
S14 Temperature-dependent CEST profiles and van't Hoff plots – T5H6 . . . . . S15
S15 Temperature-dependent CEST profiles and van't Hoff plots – A8H2 . . . . . S16
S16 Temperature-dependent CEST profiles and van't Hoff plots – A8H8 . . . . . S17
S17 Temperature-dependent CEST profiles and van't Hoff plots – T9H6 . . . . . S18
S18 Temperature-dependent CEST profiles and van't Hoff plots – C12H6 . . . . S19
S19 Concentration-dependent UV/Vis melting profiles . . . . . . . . . . . . . . . . S19
S20 $R_{1\rho}$ relaxation dispersion profiles – $C_{7.0}$ . . . . . . . . . . . . . . . . . . . . . S20
S21 $R_{1\rho}$ relaxation dispersion profiles – $caC_{7.0}$ . . . . . . . . . . . . . . . . . . . . S21
S22 $R_{1\rho}$ relaxation dispersion profiles – $caC_{5.8}$ . . . . . . . . . . . . . . . . . . . . S22
S23 $R_{1\rho}$ relaxation dispersion profiles – $caC_{4.7}$ . . . . . . . . . . . . . . . . . . . . S23
S24 $R_{1\rho}$ relaxation dispersion profiles – $fC_{7.0}$ . . . . . . . . . . . . . . . . . . . . S24

### List of Tables

S1 CEST-derived thermodynamic and kinetic parameters . . . . . . . . . . . . . . S25

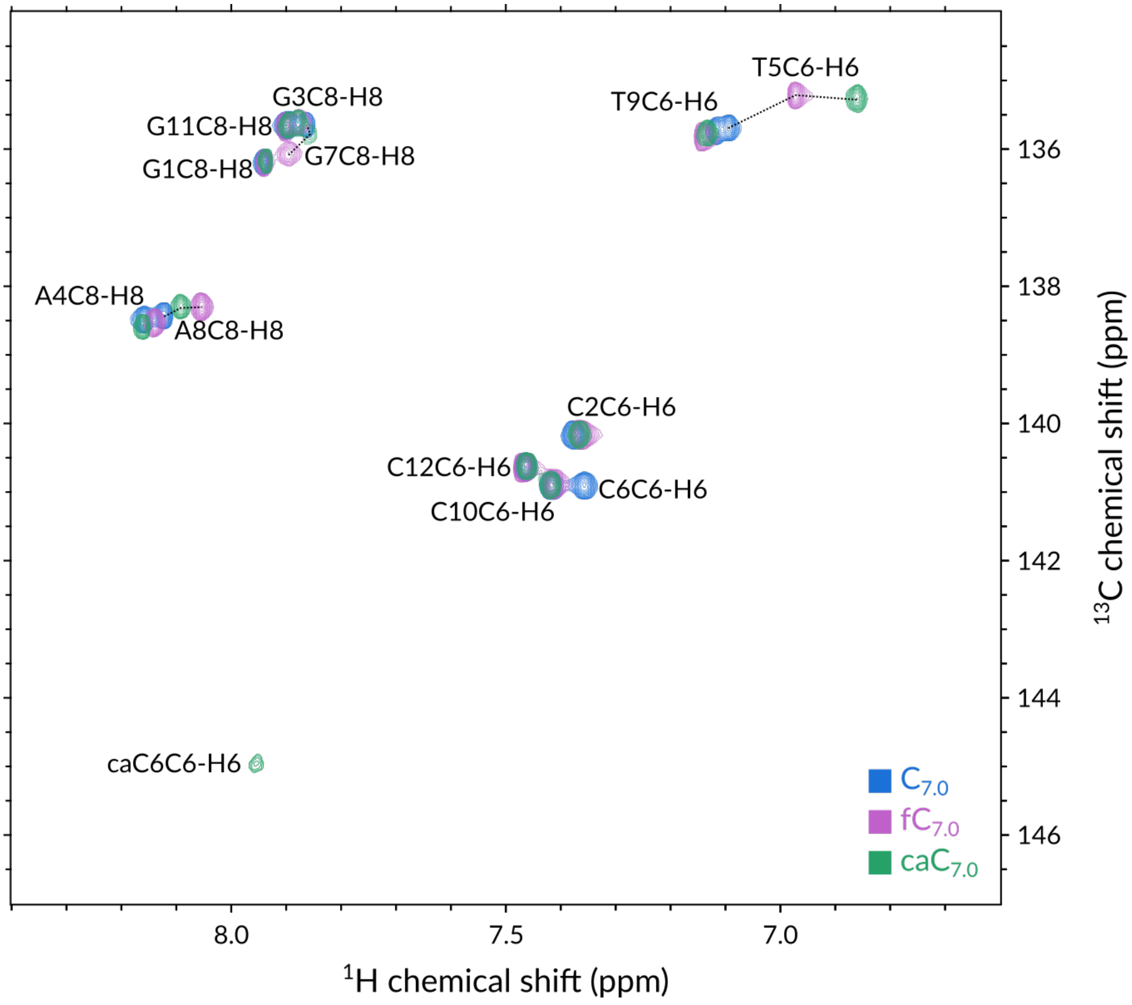


Figure S1: $^{1}$H-$^{13}$C HSQC aromatic region spectrum of $fC_{7.0}$, $caC_{7.0}$ and $C_{7.0}$ recorded at 800 MHz at 37 ℃.

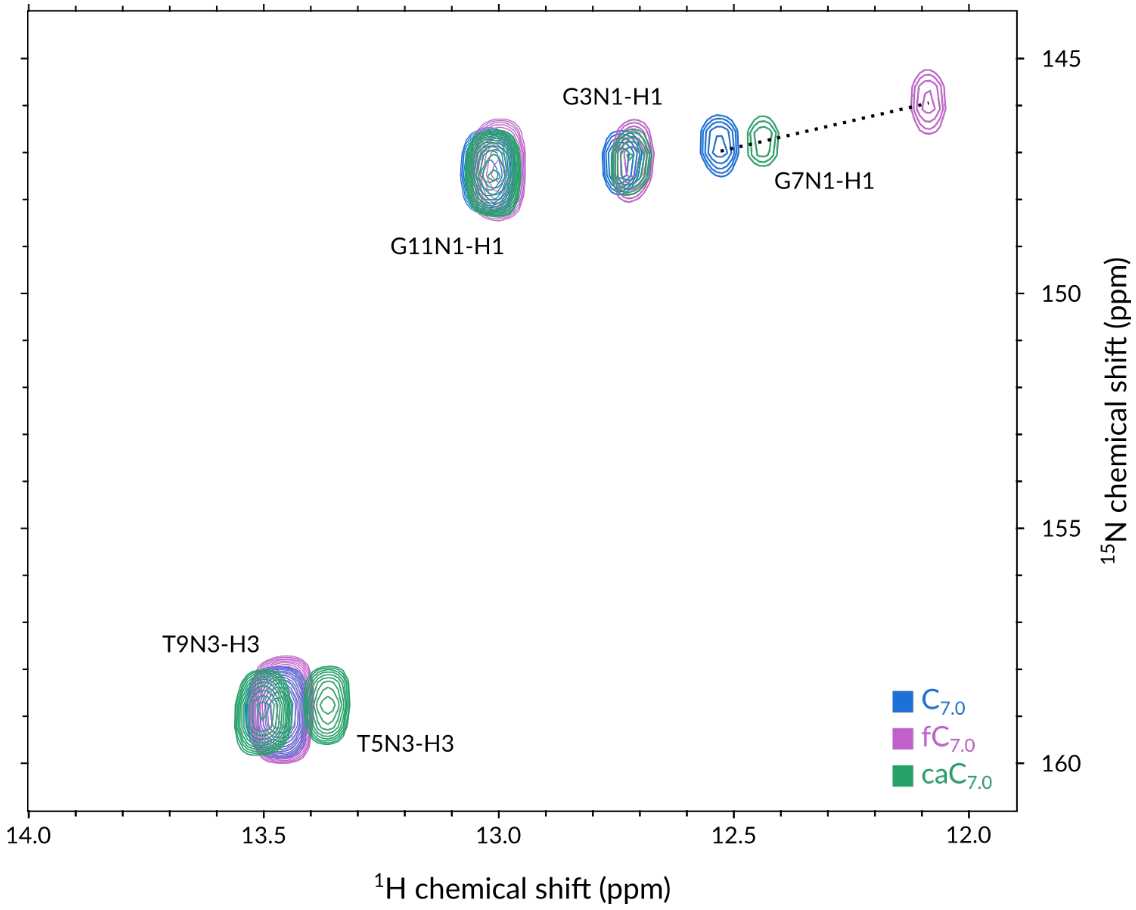


Figure S2: $^{1}$H-$^{15}$N HMQC imino region spectrum of $fC_{7.0}$, $caC_{7.0}$ and $C_{7.0}$ recorded at 800 MHz at 37 ℃.

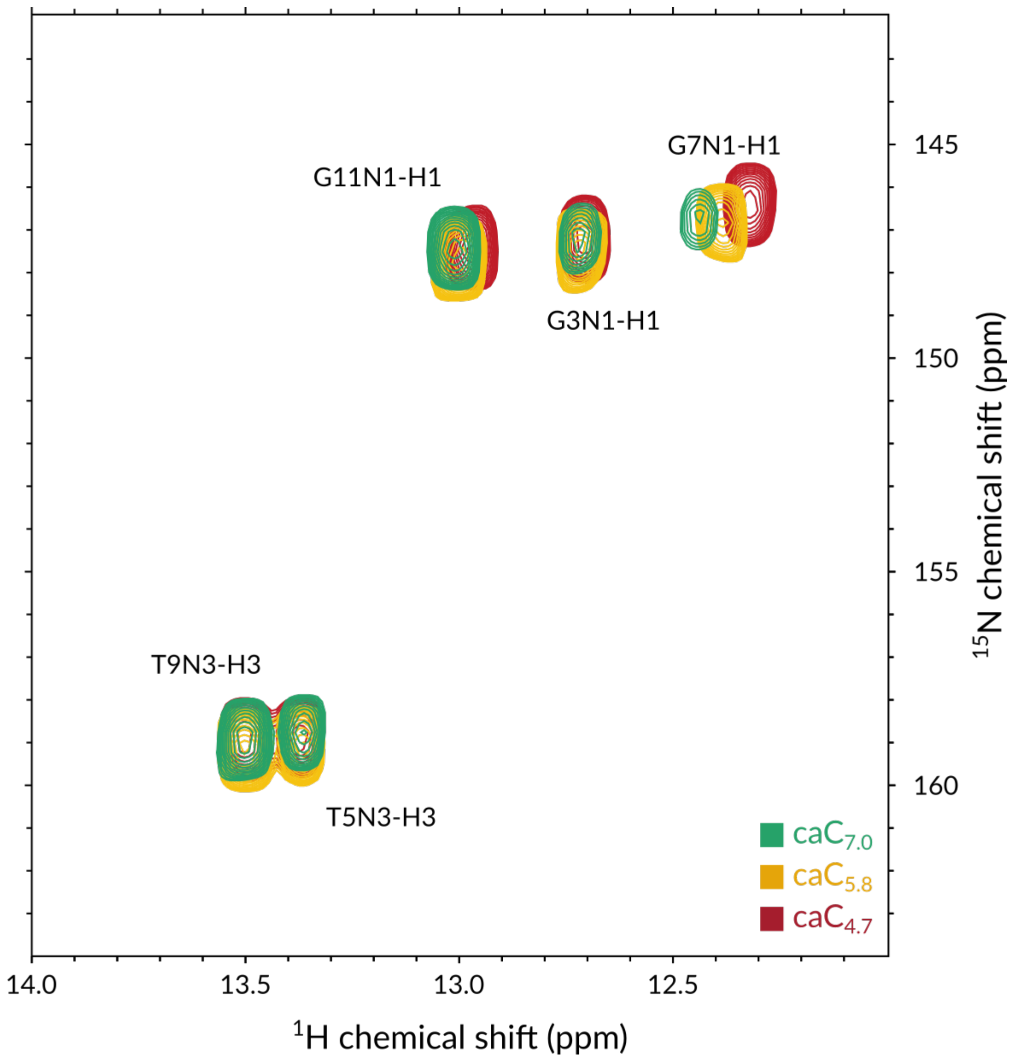


Figure S3: $^{1}$H-$^{15}$N HMQC imino region spectrum of $caC_{7.0}$, $caC_{5.8}$ and $caC_{4.7}$ recorded at 800 MHz at 37 ℃.

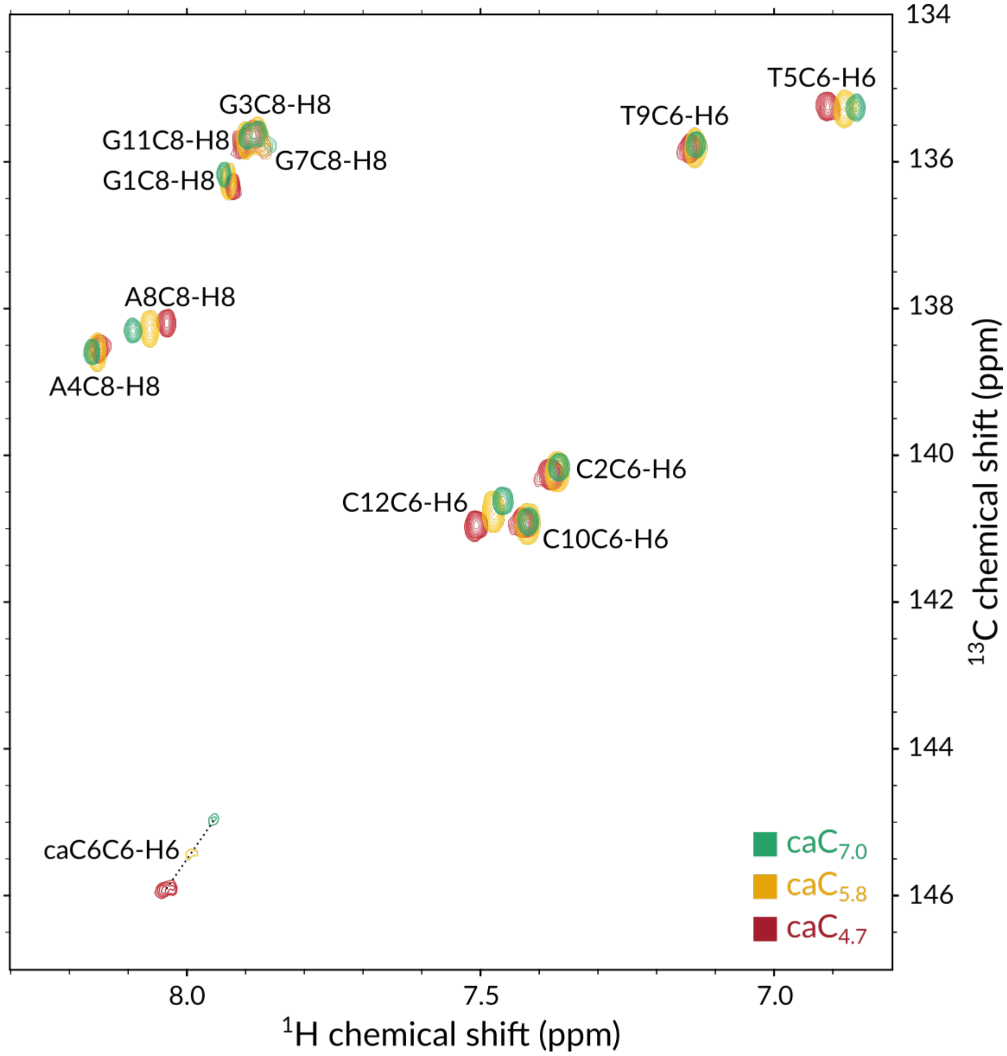


Figure S4: $^{1}$H-$^{13}$C HSQC aromatic region spectrum of caC$_{7.0}$, caC$_{5.8}$ and caC$_{4.7}$ recorded at 800 MHz at 37 °C.

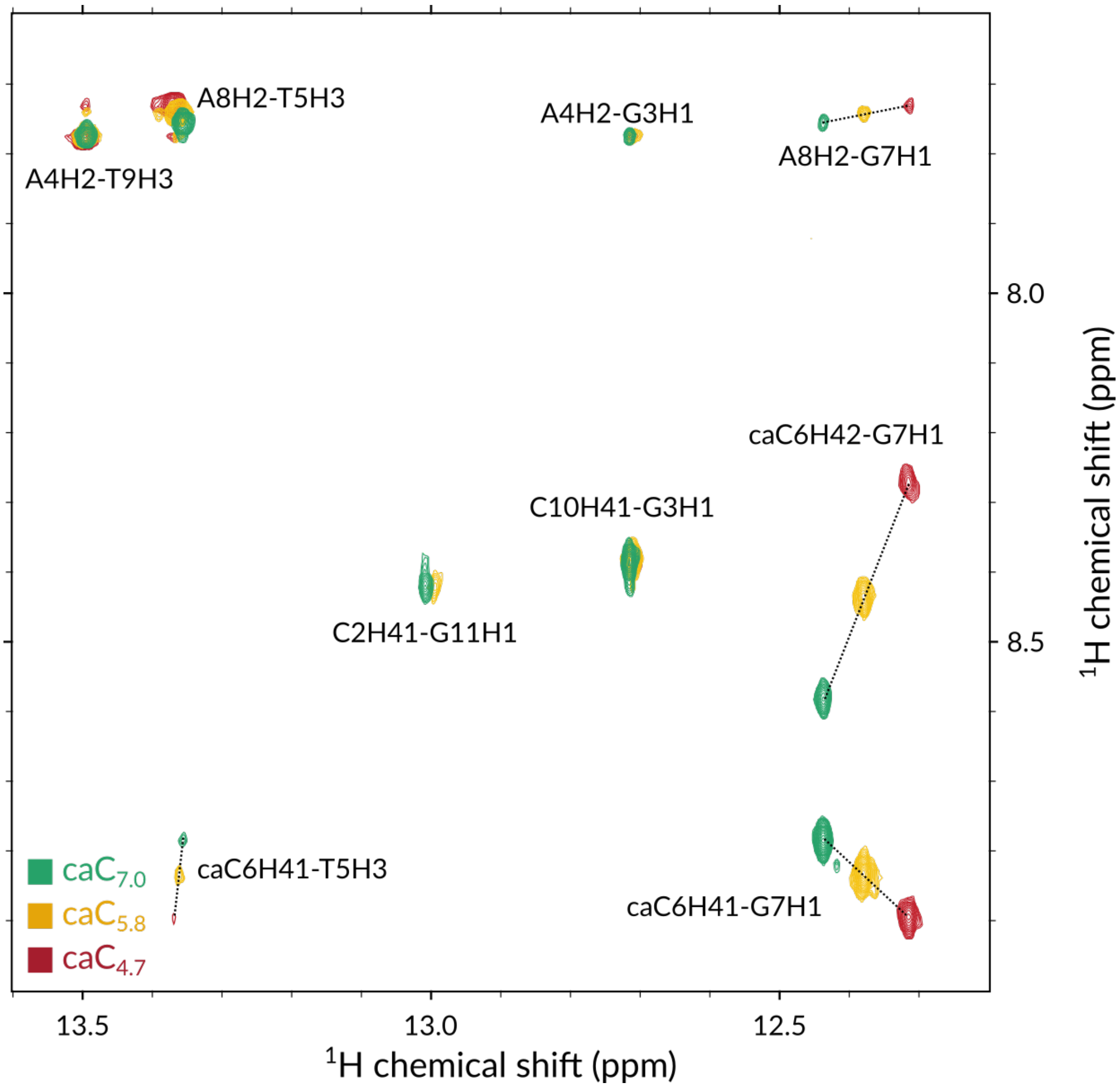


Figure S5: $^1$H-$^1$H NOESY imino region spectrum of caC$_{7.0}$, caC$_{5.8}$ and caC$_{4.7}$ recorded at 800 MHz at 37 ℃.

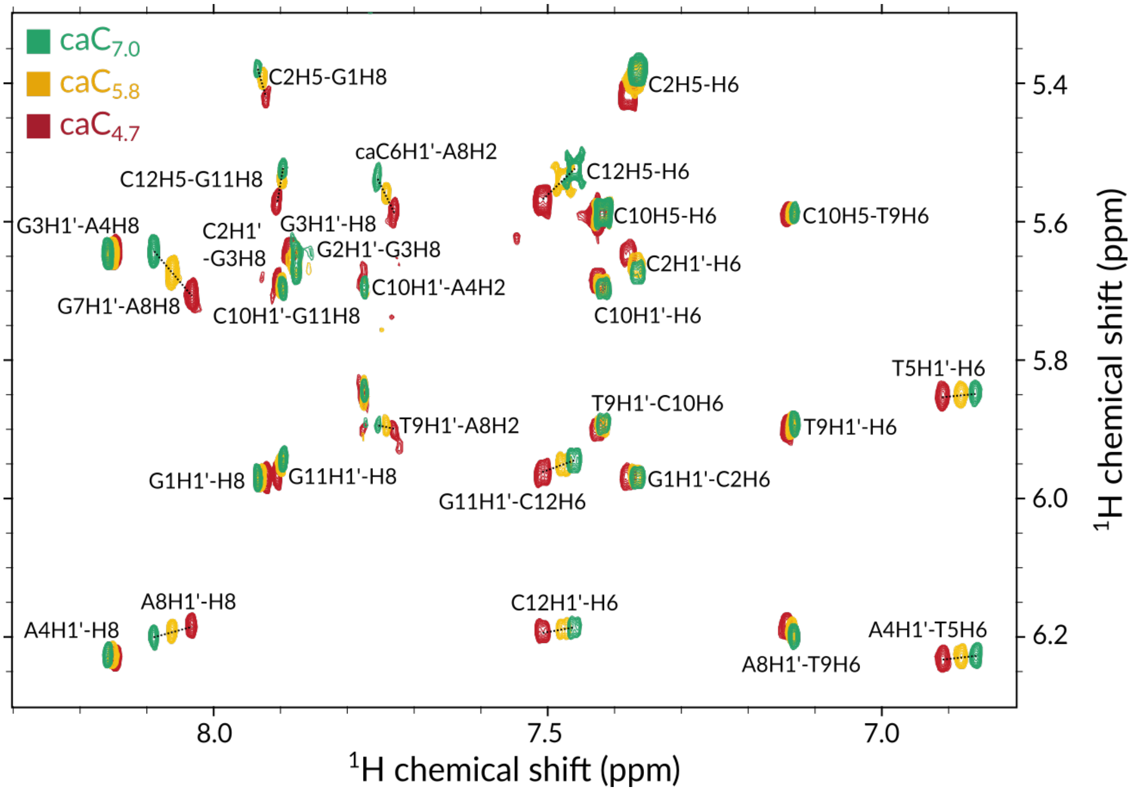


Figure S6: $^{1}$H-$^{1}$H NOESY aromatic region spectrum of caC$_{7.0}$, caC$_{5.8}$ and caC$_{4.7}$ recorded at 800 MHz at 37 ℃.

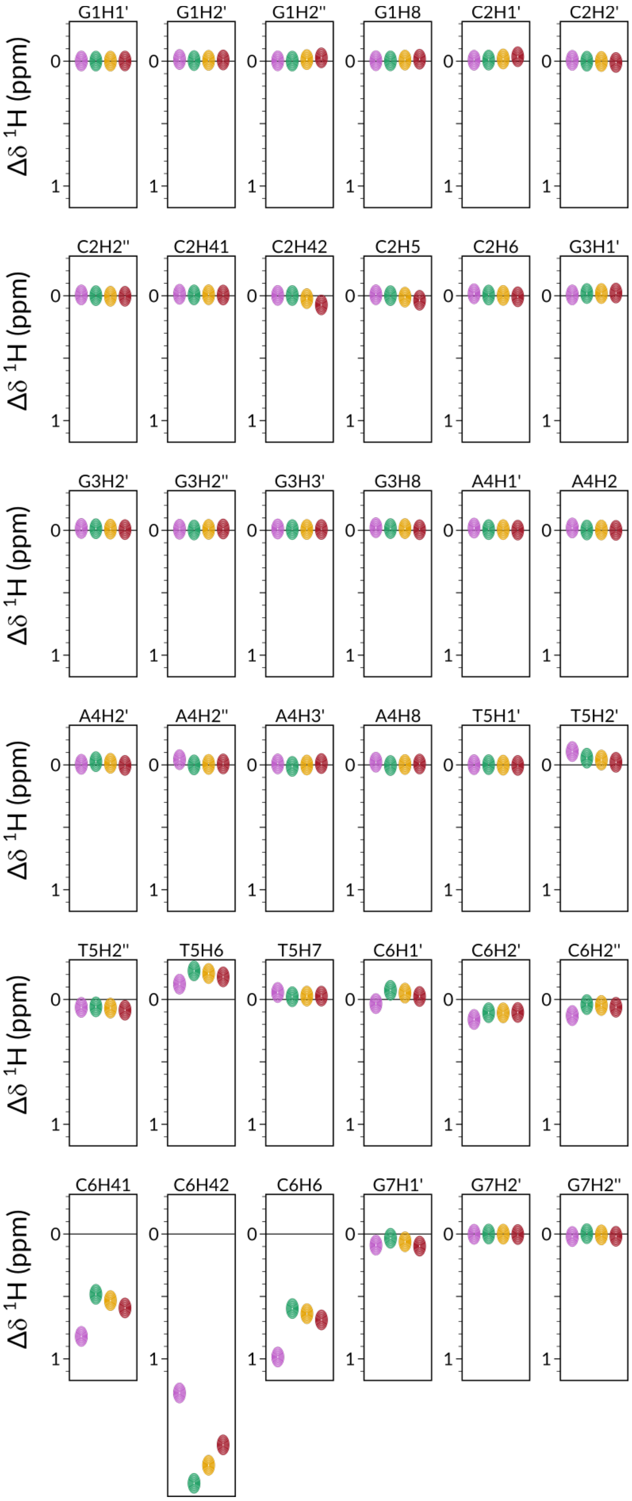


Figure S7: Comparison of chemical shift perturbations $\Delta\delta$ for all comparable aliphatic and aromatic $^1$H nuclei across all samples in identical conditions. Resonances are reported with respect to the chemical shifts of $C_{7.0}$. Magenta, green, yellow and red symbols represent $fC_{7.0}$, $caC_{7.0}$, $caC_{5.8}$ and $caC_{4.7}$, respectively. Remainder of the figure follows in Fig. S3.

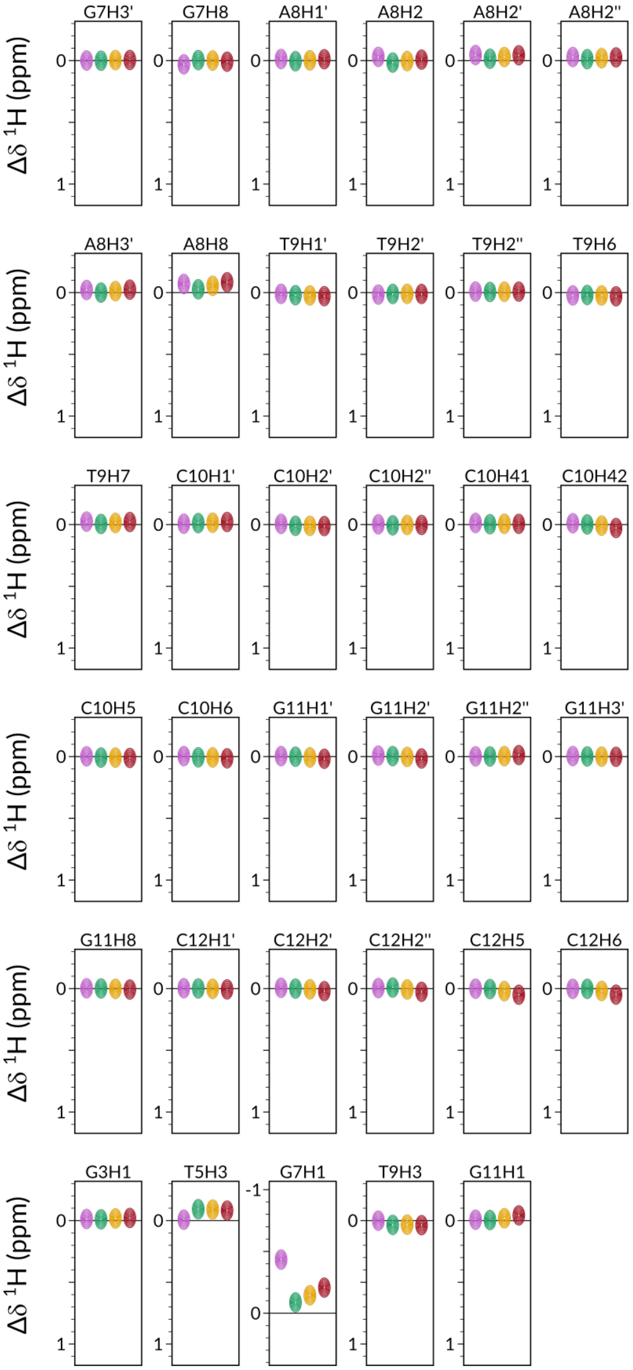


Figure S8: Comparison of chemical shift perturbations $\Delta\delta$ for all comparable aliphatic and aromatic $^1$H nuclei across all samples in identical conditions. Resonances are reported with respect to the chemical shifts of $C_{7.0}$. Magenta, green, yellow and red symbols represent $fC_{7.0}$, $caC_{7.0}$, $caC_{5.8}$ and $caC_{4.7}$, respectively.

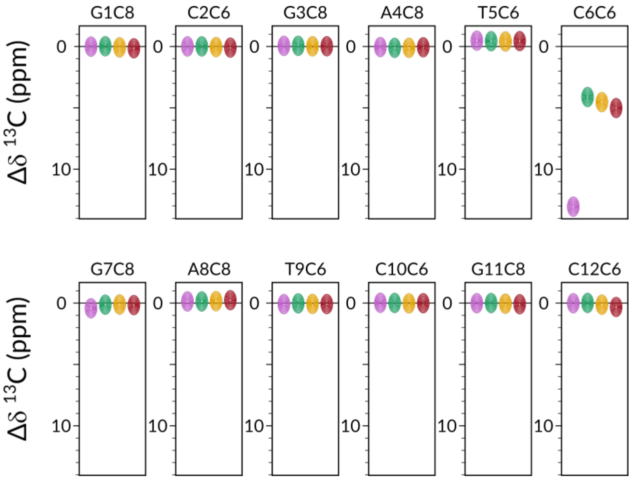


Figure S9: Comparison of chemical shift perturbations $\Delta\delta$ for all comparable aromatic $^{13}C$ nuclei across all samples in identical conditions. Resonances are reported with respect to the chemical shifts of $C_{7.0}$. Magenta, green, yellow and red symbols represent $fC_{7.0}$, $caC_{7.0}$, $caC_{5.8}$ and $caC_{4.7}$, respectively.

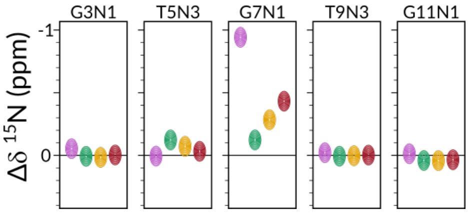


Figure S10: Comparison of chemical shift perturbations $\Delta\delta$ for all comparable aromatic $^{15}N$ nuclei across all samples in identical conditions. Resonances are reported with respect to the chemical shifts of $C_{7.0}$. Magenta, green, yellow and red symbols represent $fC_{7.0}$, $caC_{7.0}$, $caC_{5.8}$ and $caC_{4.7}$, respectively.

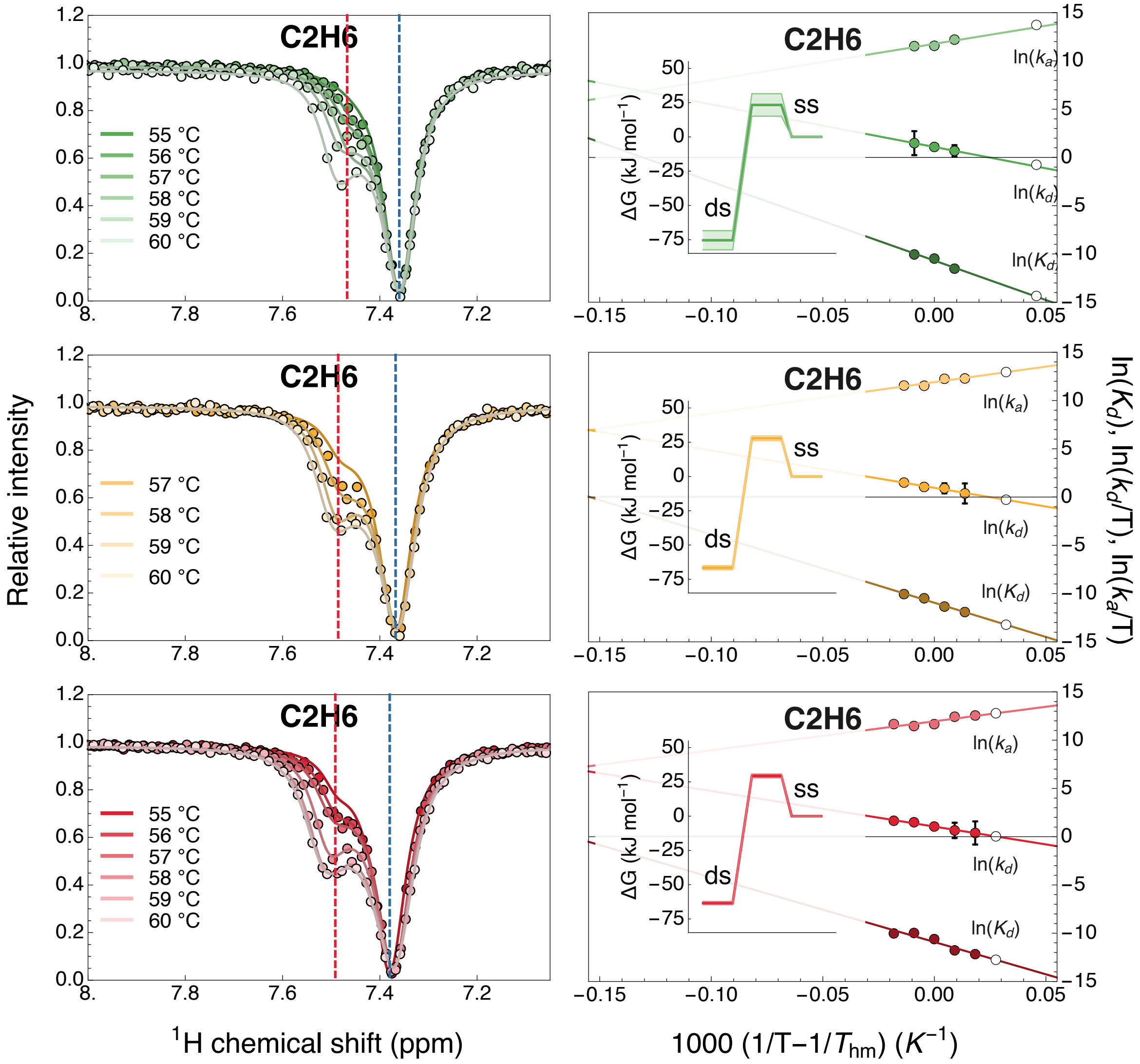


Figure S11: Temperature-dependent CEST melting profiles and van't Hoff plots for proton C2H6. $caC_{7.0}$, $caC_{5.8}$ and $caC_{4.7}$ are shown in shades of green, yellow and red, respectively. Dashed blue and red lines indicate GS and ES chemical shift values. In the van't Hoff plots, shades of green, yellow and red indicate data entries and linear fits for $caC_{7.0}$, $caC_{5.8}$ and $caC_{4.7}$, respectively. White data points represent back-calculated values. Insets present the relevant Gibbs free energy plots at each pH condition at 37 °C.

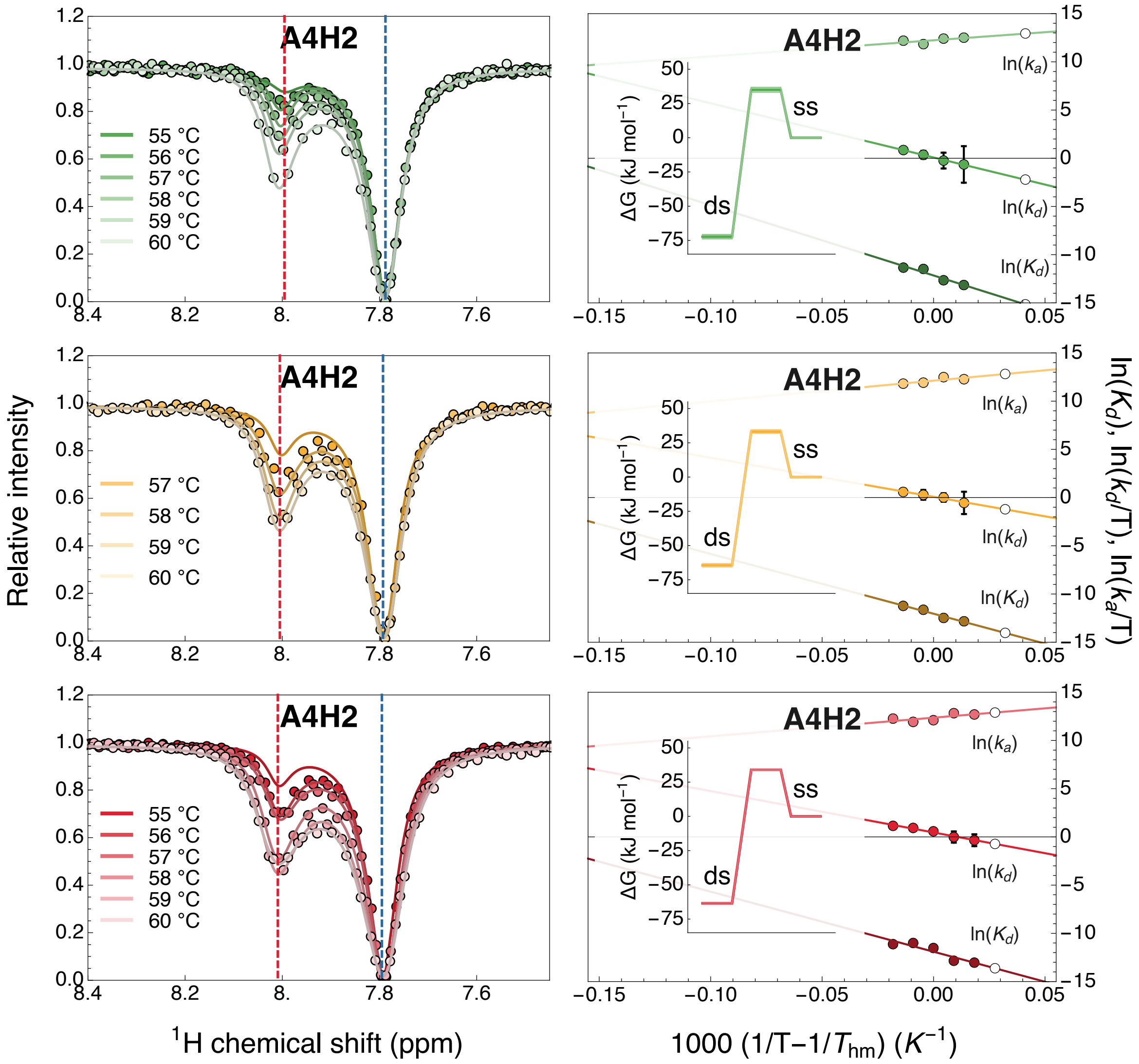


Figure S12: Temperature-dependent CEST melting profiles and van't Hoff plots for proton A4H2. $caC_{7.0}$, $caC_{5.8}$ and $caC_{4.7}$ are shown in shades of green, yellow and red, respectively. Dashed blue and red lines indicate GS and ES chemical shift values. In the van't Hoff plots, shades of green, yellow and red indicate data entries and linear fits for $caC_{7.0}$, $caC_{5.8}$ and $caC_{4.7}$, respectively. White data points represent back-calculated values. Insets present the relevant Gibbs free energy plots at each pH condition at 37 °C.

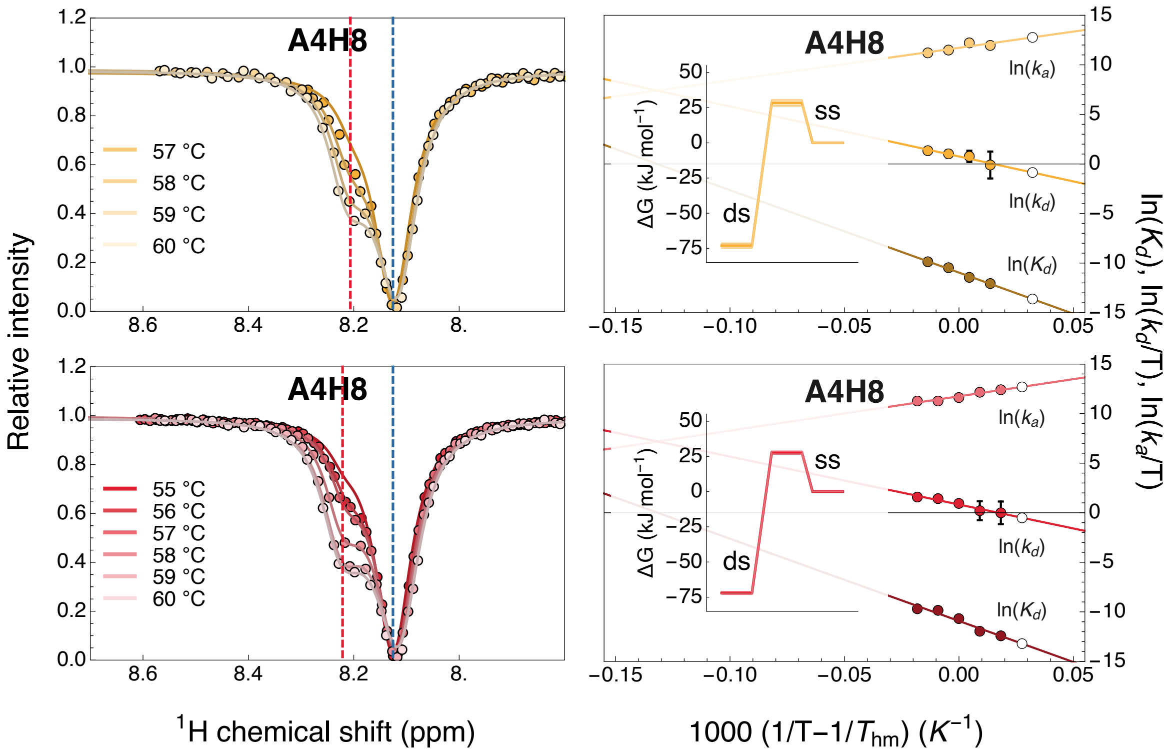


Figure S13: Temperature-dependent CEST melting profiles and van't Hoff plots for proton A4H8. $caC_{5.8}$ and $caC_{4.7}$ are shown in shades of yellow and red, respectively. Dashed blue and red lines indicate GS and ES chemical shift values. In the van't Hoff plots, shades of yellow and red indicate data entries and linear fits for $caC_{5.8}$ and $caC_{4.7}$, respectively. White data points represent back-calculated values. Insets present the relevant Gibbs free energy plots at each pH condition at 37 °C.

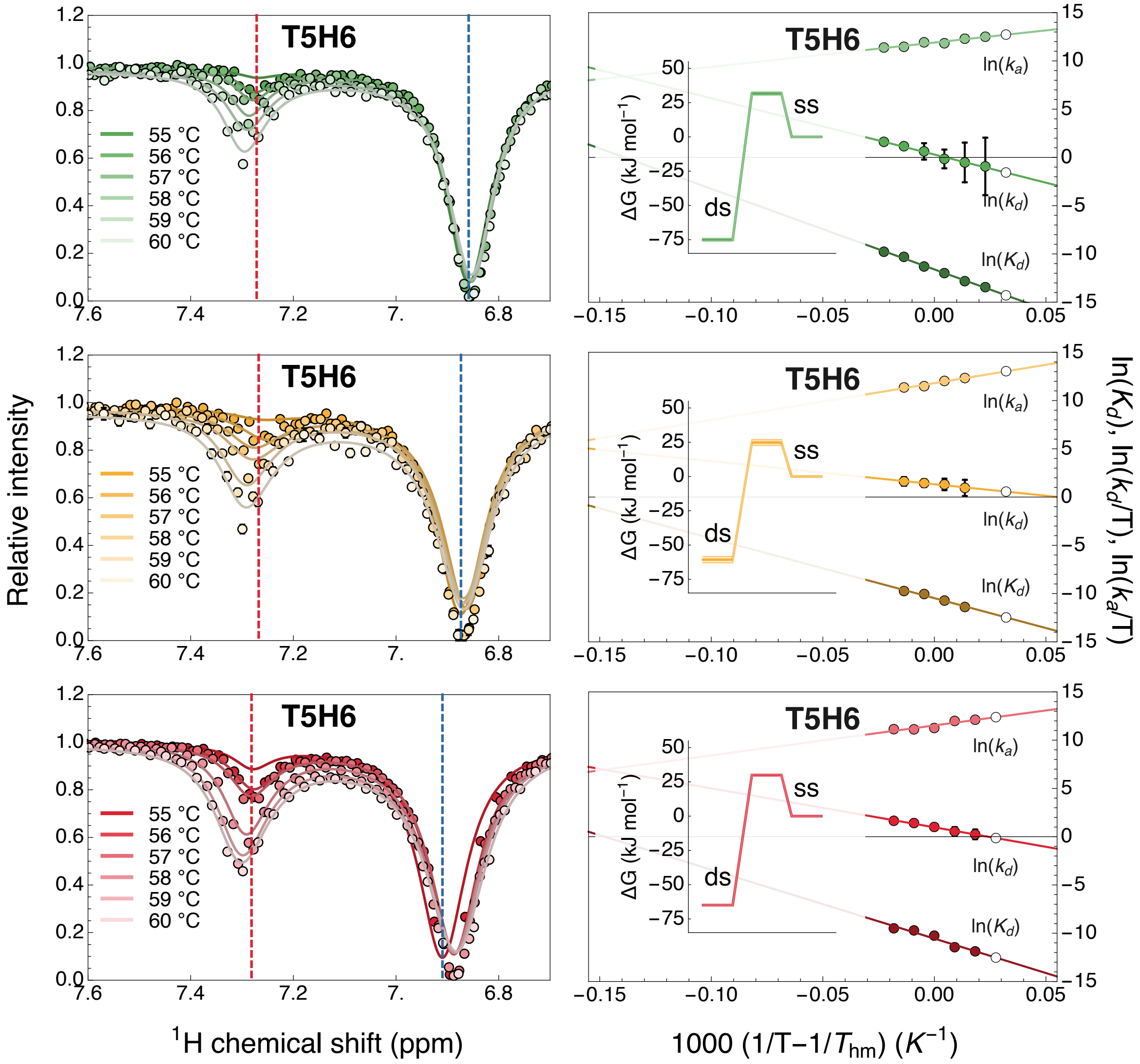


Figure S14: Temperature-dependent CEST melting profiles and van't Hoff plots for proton T5H6. $caC_{7.0}$, $caC_{5.8}$ and $caC_{4.7}$ are shown in shades of green, yellow and red, respectively. Dashed blue and red lines indicate GS and ES chemical shift values. In the van't Hoff plots, shades of green, yellow and red indicate data entries and linear fits for $caC_{7.0}$, $caC_{5.8}$ and $caC_{4.7}$, respectively. White data points represent back-calculated values. Insets present the relevant Gibbs free energy plots at each pH condition at 37 °C.

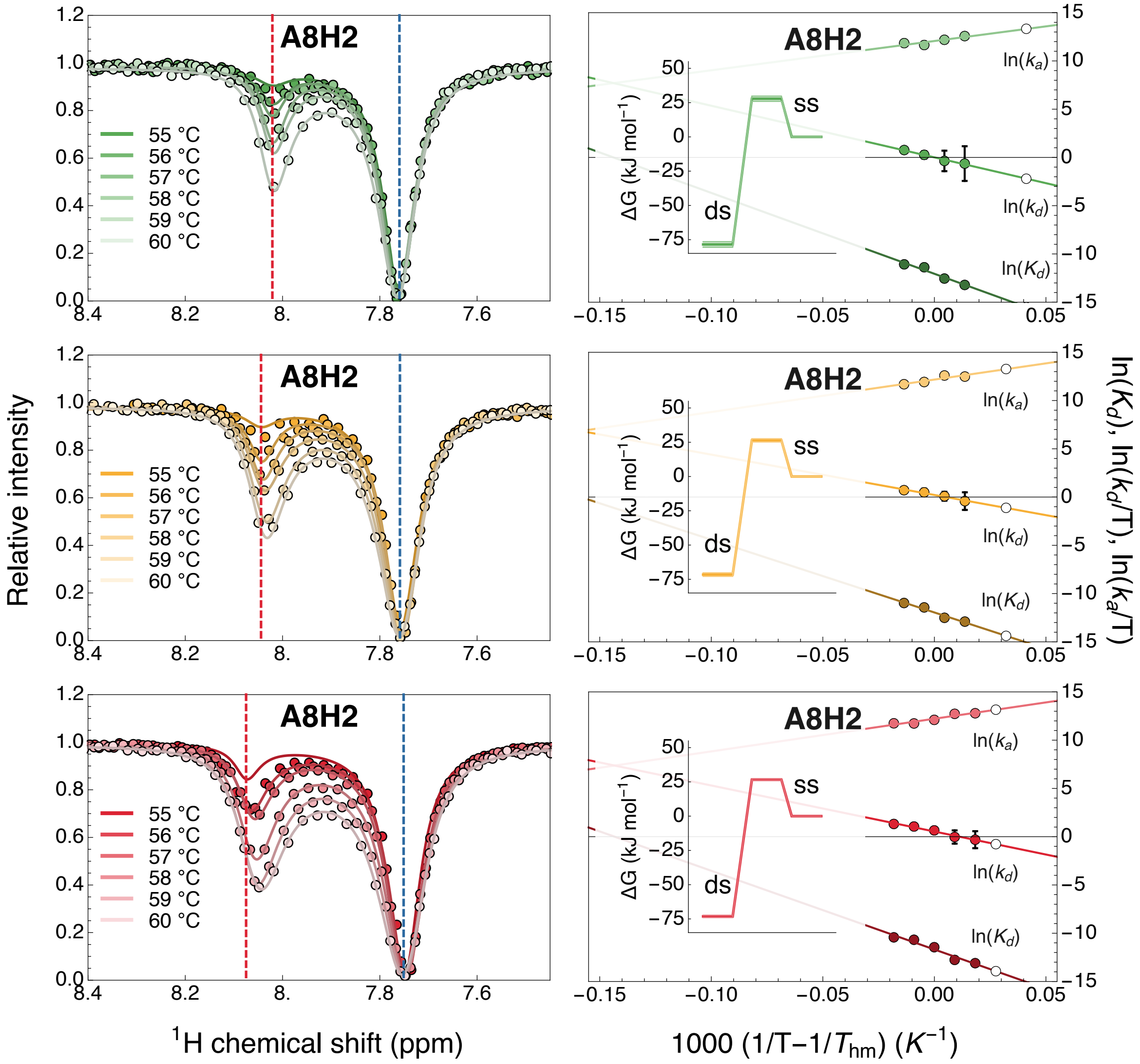


Figure S15: Temperature-dependent CEST melting profiles and van't Hoff plots for proton A8H2. $caC_{7.0}$, $caC_{5.8}$ and $caC_{4.7}$ are shown in shades of green, yellow and red, respectively. Dashed blue and red lines indicate GS and ES chemical shift values. In the van't Hoff plots, shades of green, yellow and red indicate data entries and linear fits for $caC_{7.0}$, $caC_{5.8}$ and $caC_{4.7}$, respectively. White data points represent back-calculated values. Insets present the relevant Gibbs free energy plots at each pH condition at 37 °C.

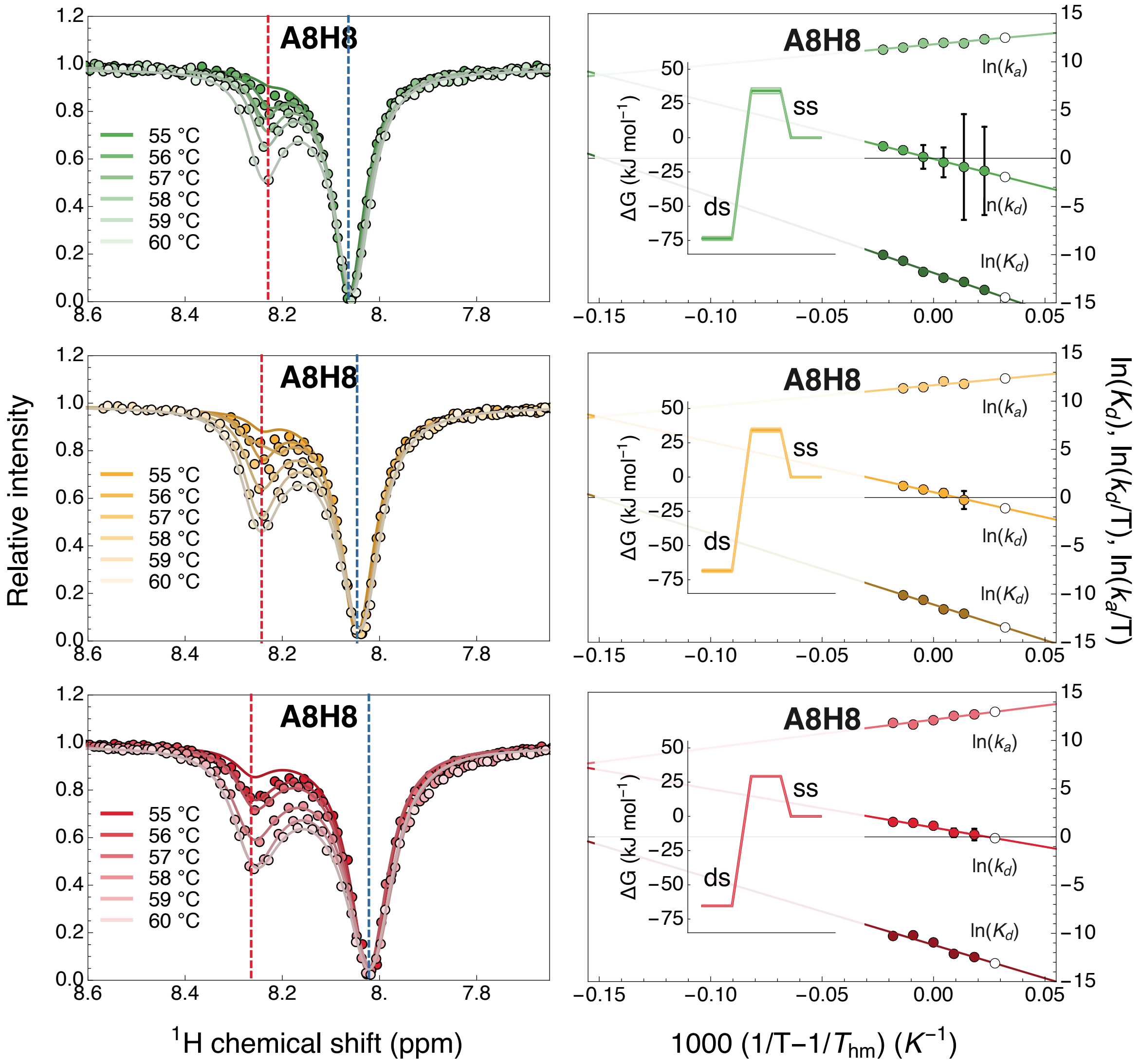


Figure S16: Temperature-dependent CEST melting profiles and van't Hoff plots for proton A8H8. $caC_{7.0}$, $caC_{5.8}$ and $caC_{4.7}$ are shown in shades of green, yellow and red, respectively. Dashed blue and red lines indicate GS and ES chemical shift values. In the van't Hoff plots, shades of green, yellow and red indicate data entries and linear fits for $caC_{7.0}$, $caC_{5.8}$ and $caC_{4.7}$, respectively. White data points represent back-calculated values. Insets present the relevant Gibbs free energy plots at each pH condition at 37 °C.

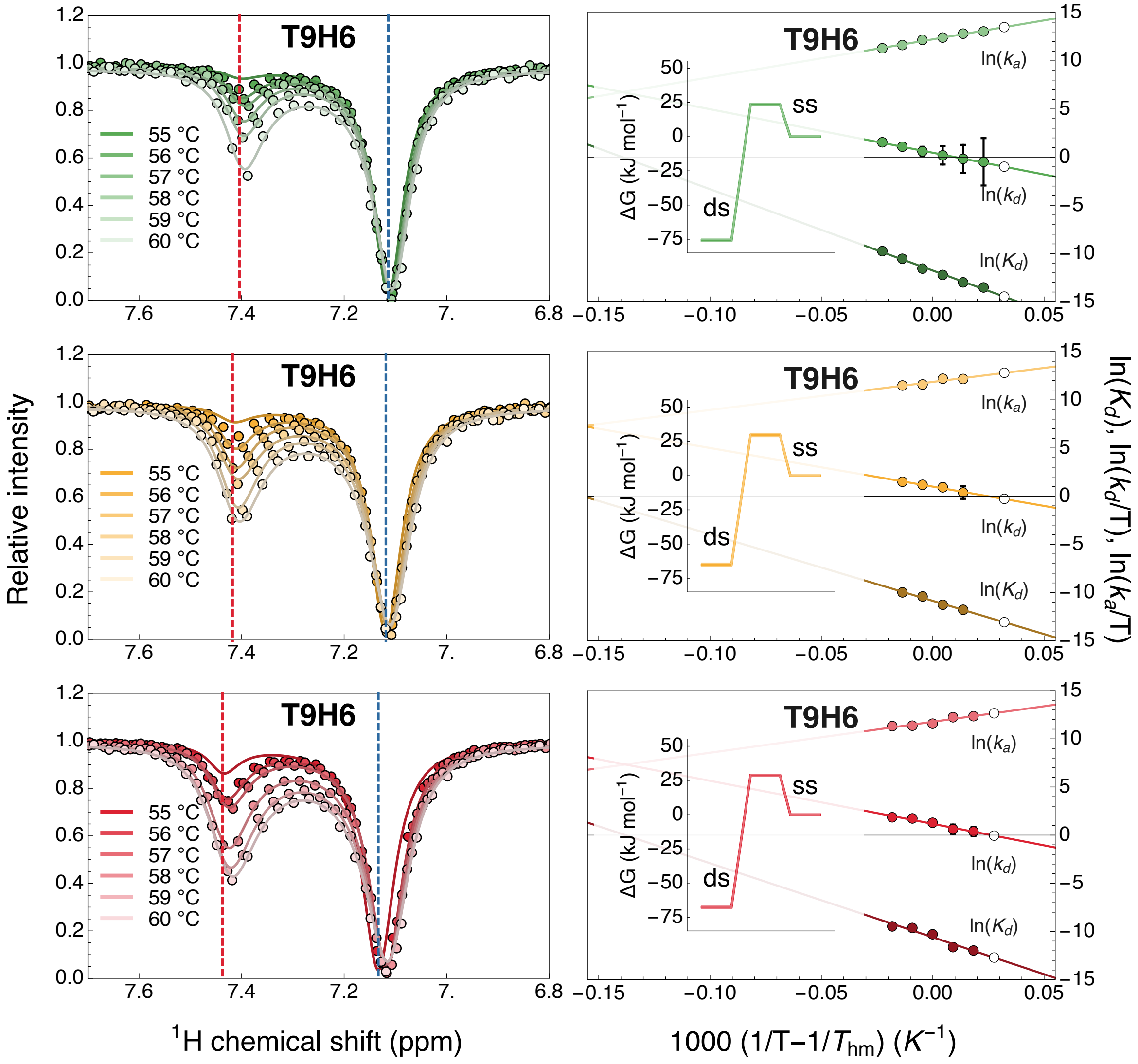


Figure S17: Temperature-dependent CEST melting profiles and van't Hoff plots for proton T9H6. $caC_{7.0}$, $caC_{5.8}$ and $caC_{4.7}$ are shown in shades of green, yellow and red, respectively. Dashed blue and red lines indicate GS and ES chemical shift values. In the van't Hoff plots, shades of green, yellow and red indicate data entries and linear fits for $caC_{7.0}$, $caC_{5.8}$ and $caC_{4.7}$, respectively. White data points represent back-calculated values. Insets present the relevant Gibbs free energy plots at each pH condition at 37 °C.

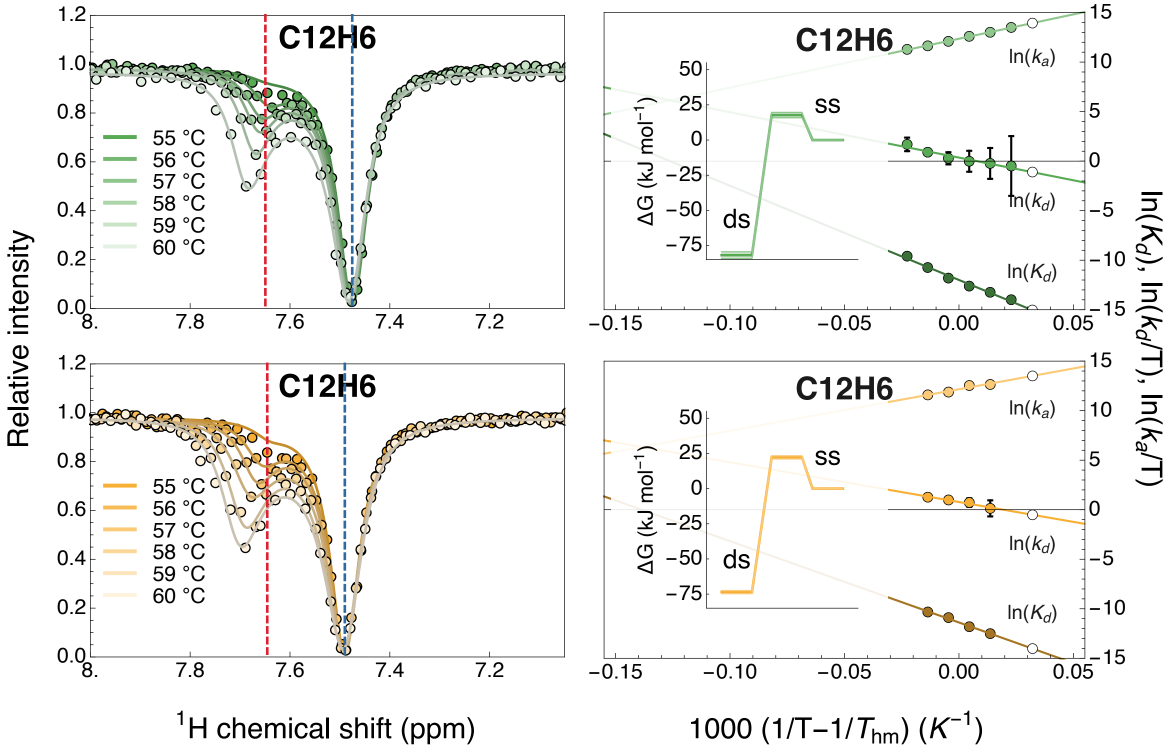


Figure S18: Temperature-dependent CEST melting profiles and van't Hoff plots for proton C12H6. $caC_{7.0}$, and $caC_{5.8}$ are shown in shades of green and yellow, respectively. Dashed blue and red lines indicate GS and ES chemical shift values. In the van't Hoff plots, shades of green and yellow indicate data entries and linear fits for $caC_{7.0}$ and $caC_{5.8}$, respectively. White data points represent back-calculated values. Insets present the relevant Gibbs free energy plots at each pH condition at 37 °C.

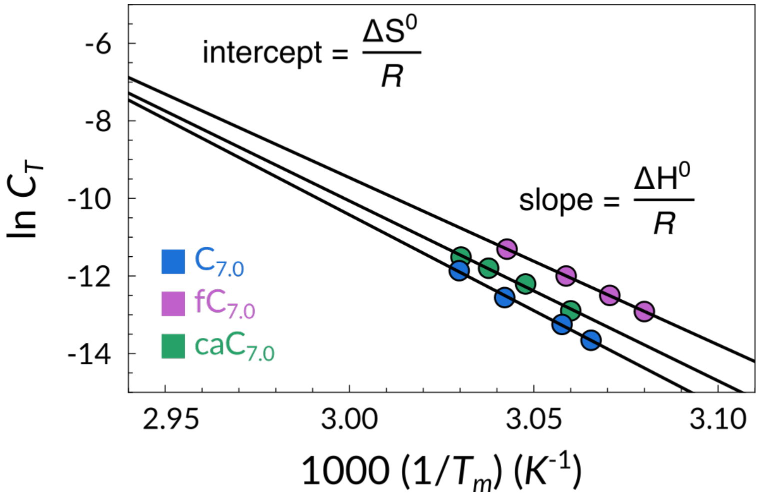


Figure S19: Concentration-dependent UV/Vis melting temperatures analysis of $C_{7.0}$ (blue), $fC_{7.0}$ (magenta) and $caC_{7.0}$ (green). The reciprocal of the observed $T_m$ is plotted as a function of the logarithm of the total DNA concentration ($C_t$). The slope and the intercept of the fitted linear corresponds to $\Delta H^0/R$, and $\Delta S^0/R$, respectively.

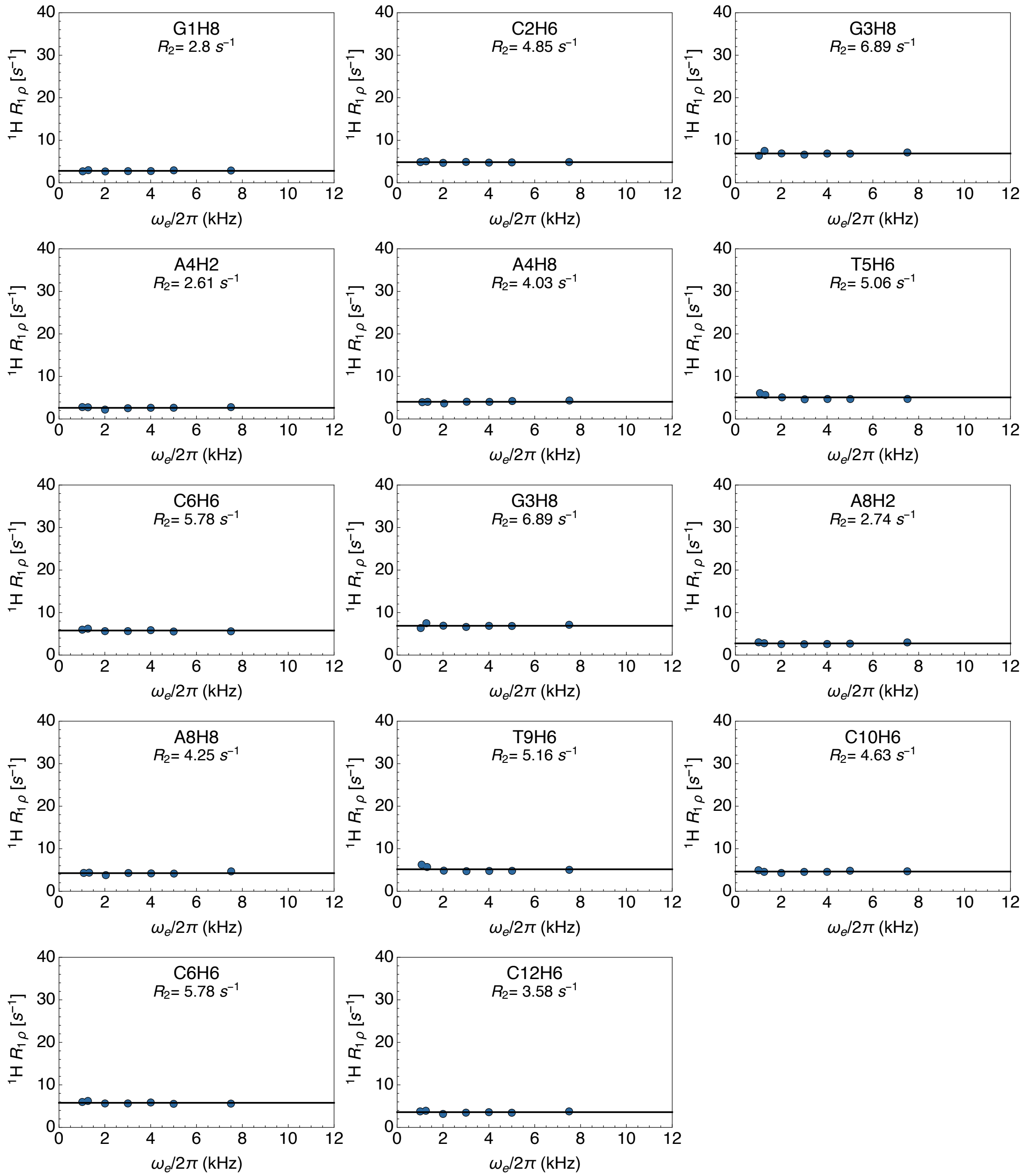


Figure S20: $R_{1\rho}$ relaxation dispersion profiles for each available proton reported of $C_{7.0}$ recorded at 55 °C. Lines represent best fits either to models accounting for (color-coded) or discounting chemical exchange (black). Fitted $R_2$ relaxation rates are displayed for each site.

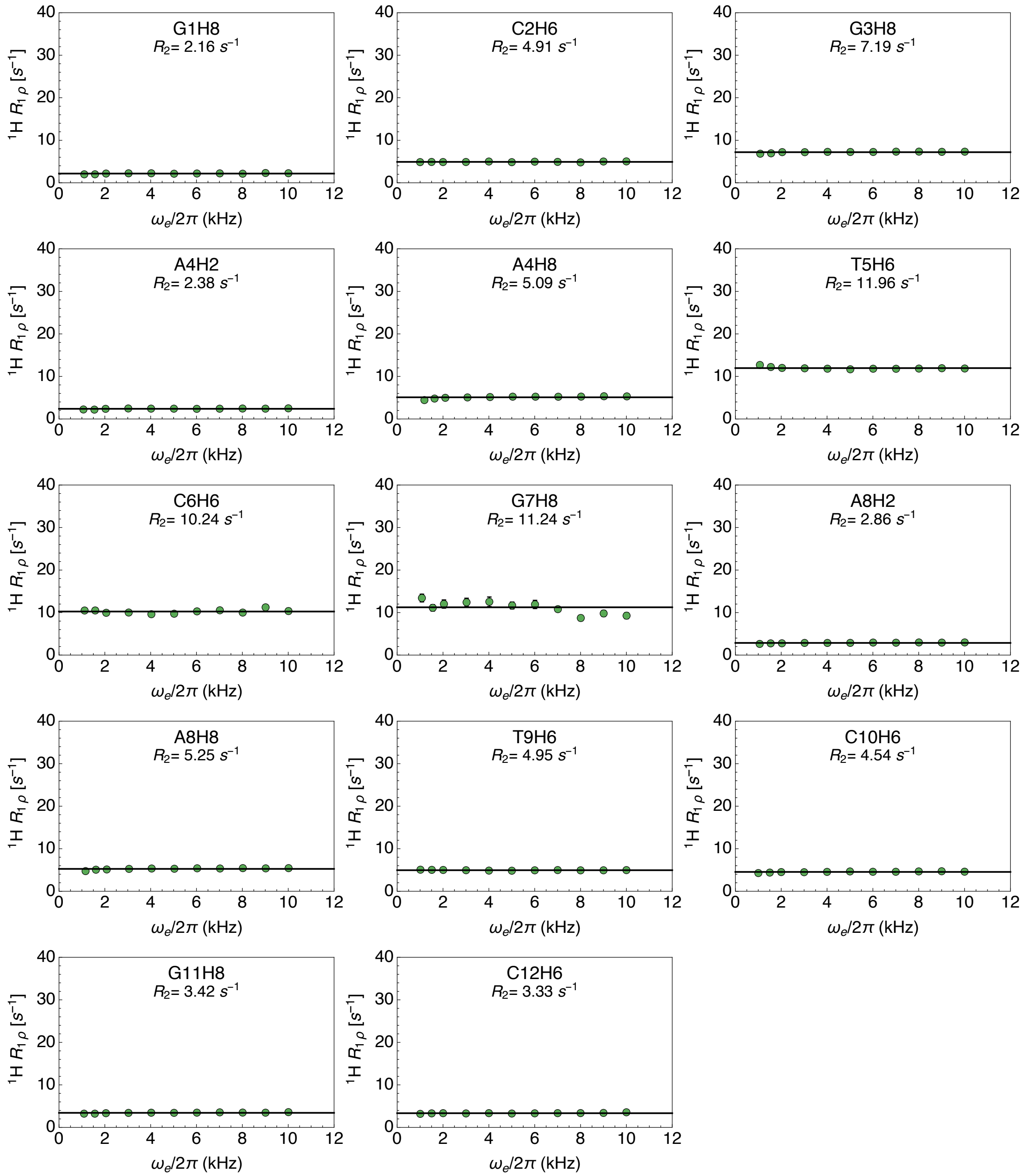


Figure S21: $R_{1\rho}$ relaxation dispersion profiles for each available proton reported of $caC_{7.0}$ recorded at 55 °C. Lines represent best fits either to models accounting for (color-coded) or discounting chemical exchange (black). Fitted $R_2$ relaxation rates are displayed for each site.

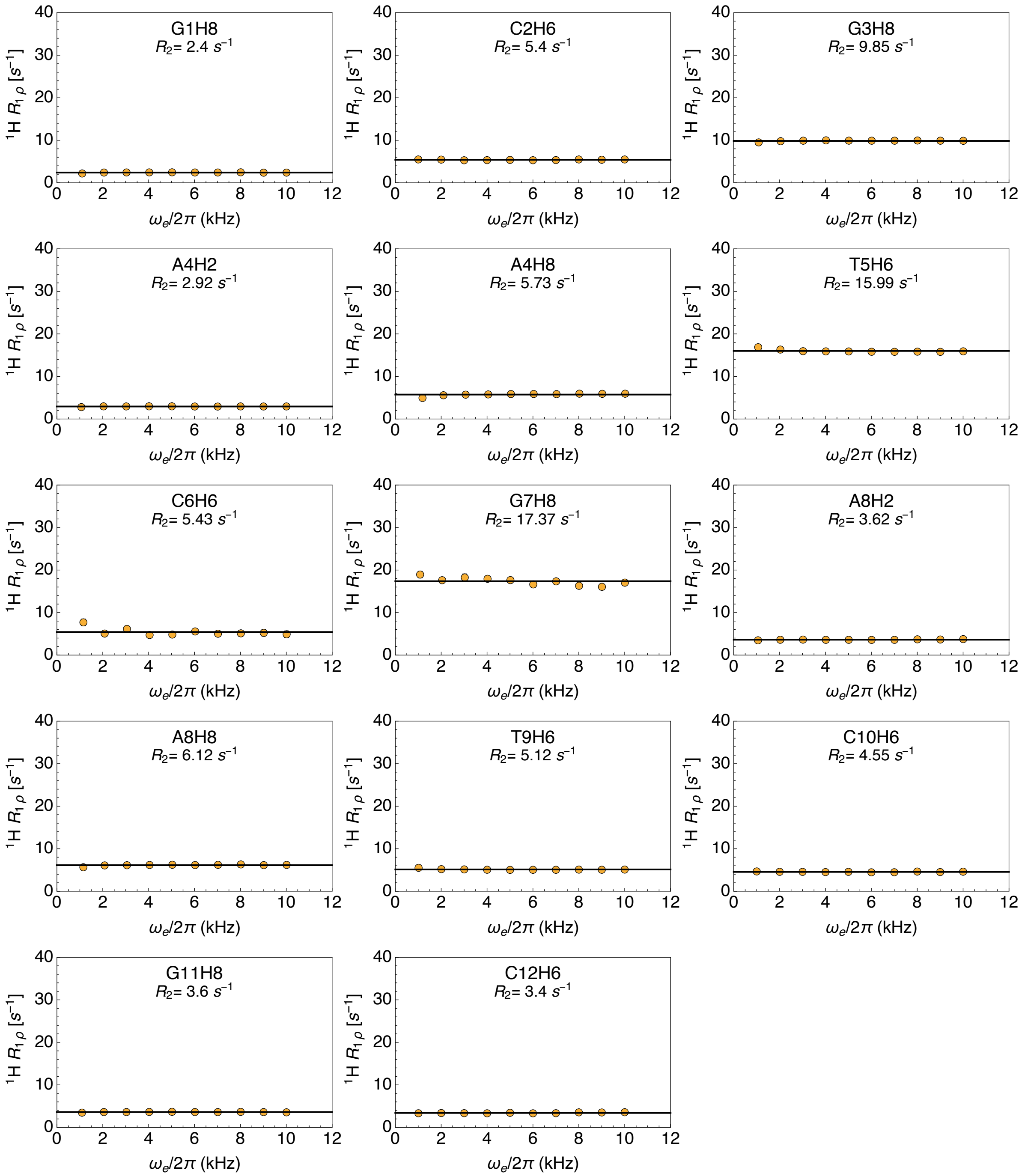


Figure S22: $R_{1\rho}$ relaxation dispersion profiles for each available proton reported of $caC_{5.8}$ recorded at 55 ℃. Lines represent best fits either to models accounting for (color-coded) or discounting chemical exchange (black). Fitted $R_2$ relaxation rates are displayed for each site.

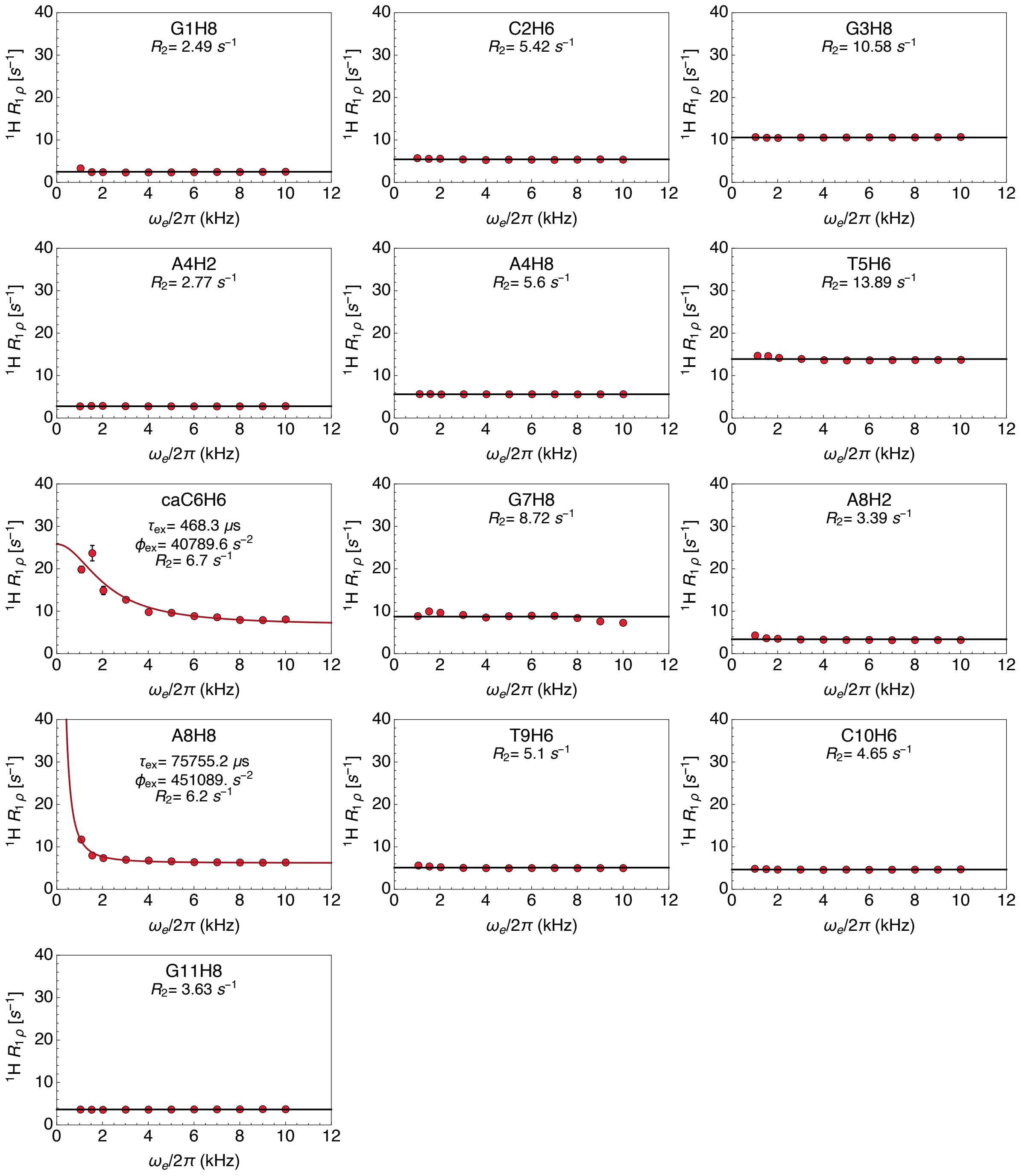


Figure S23: $R_{1\rho}$ relaxation dispersion profiles for each available proton reported of $caC_{4.7}$ recorded at 55 ℃. Lines represent best fits either to models accounting for (color-coded) or discounting chemical exchange (black). Fitted $R_2$ relaxation rates are displayed for each site, while exchange parameters such as $\tau_{ex}$ and $\phi_{ex}$ are only available for profiles best fit to the exchange model.

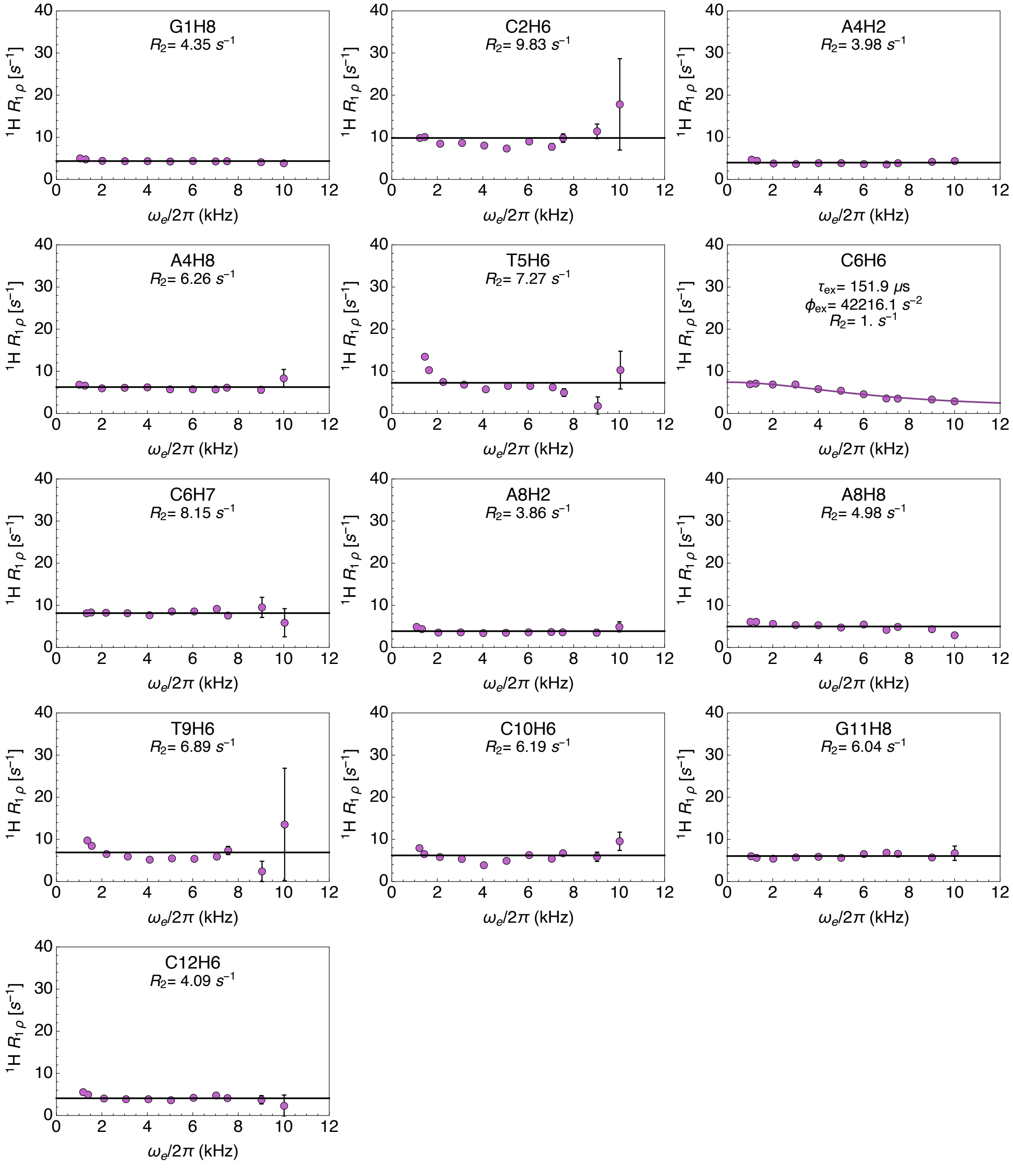


Figure S24: $R_{1\rho}$ relaxation dispersion profiles for each available proton reported of $fC_{7.0}$ recorded at 55 °C. Lines represent best fits either to models accounting for (color-coded) or discounting chemical exchange (black). Fitted $R_2$ relaxation rates are displayed for each site, while exchange parameters such as $\tau_{ex}$ and $\phi_{ex}$ are only available for profiles best fit to the exchange model.

Table S1: Thermodynamic and kinetic parameters of the dsDNA melting process obtained from the van't Hoff and Eyring analysis of the CEST-derived exchange parameters. Errors are given as one standard deviation.

| | | **Thermodynamics** | | | | **Dissociation kinetics** | | | **Association kinetics** | | |
|---|---|---|---|---|---|---|---|---|---|---|---|
| | | $\Delta G^0_{37\mathbf{C}}$ | $\Delta H^0$ | $\Delta S^0$ | $T_m$ (1 mM) | $\Delta G^{\ddagger}_{d,37\mathbf{C}}$ | $\Delta H^{\ddagger}_d$ | $\Delta S^{\ddagger}_d$ | $\Delta G^{\ddagger}_{a,37\mathbf{C}}$ | $\Delta H^{\ddagger}_a$ | $\Delta S^{\ddagger}_a$ |
| | | kJ mol$^{-1}$ | kJ mol$^{-1}$ | J K$^{-1}$ mol$^{-1}$ | °C | kJ mol$^{-1}$ | kJ mol$^{-1}$ | J K$^{-1}$ mol$^{-1}$ | kJ mol$^{-1}$ | kJ mol$^{-1}$ | J K$^{-1}$ mol$^{-1}$ |
| $C_{7.0}$ | C2H6 | 71.1 ± 1.7 | 641 ± 26 | 1838 ± 77 | 65 ± 0.2 | 99.2 ± 2.2 | 372 ± 31 | 880 ± 94 | 28.1 ± 2.1 | -269 ± 30 | -958 ± 91 |
| | G3H8 | 69. ± 1.3 | 608 ± 20 | 1738 ± 59 | 66 ± 0.2 | 106.4 ± 1.9 | 459 ± 28 | 1138 ± 84 | 37.4 ± 1.6 | -149 ± 23 | -600 ± 71 |
| | A4H2 | 72.5 ± 0.9 | 626 ± 14 | 1784 ± 41 | 67 ± 0.2 | 100.1 ± 0.7 | 363 ± 11 | 847 ± 32 | 27.7 ± 0.7 | -263 ± 11 | -936 ± 32 |
| | A4H8 | 77.6 ± 1.1 | 732 ± 16 | 2110 ± 48 | 65 ± 0.1 | 108.3 ± 1.8 | 498 ± 25 | 1258 ± 76 | 30.7 ± 1.8 | -234 ± 25 | -852 ± 75 |
| | T5H6 | 71. ± 1.6 | 633 ± 23 | 1812 ± 70 | 65 ± 0.2 | 102.7 ± 1.4 | 415 ± 20 | 1006 ± 60 | 31.7 ± 1.3 | -218 ± 19 | -806 ± 57 |
| | C6H6 | 70.7 ± 1.4 | 638 ± 21 | 1829 ± 63 | 65 ± 0.2 | 99.8 ± 1.2 | 379 ± 17 | 900 ± 51 | 29.1 ± 1.2 | -259 ± 17 | -929 ± 51 |
| | A8H2 | 73.8 ± 1.1 | 643 ± 17 | 1834 ± 50 | 67 ± 0.2 | 103.6 ± 1. | 409 ± 15 | 986 ± 45 | 29.8 ± 1. | -233 ± 14 | -849 ± 42 |
| | A8H8 | 73.1 ± 1. | 666 ± 15 | 1910 ± 45 | 65 ± 0.1 | 104.3 ± 1. | 439 ± 14 | 1078 ± 43 | 31.2 ± 0.9 | -227 ± 13 | -833 ± 40 |
| | T9H6 | 70.2 ± 1.8 | 626 ± 27 | 1791 ± 80 | 65 ± 0.3 | 102.2 ± 1.5 | 410 ± 22 | 993 ± 66 | 31.9 ± 1.5 | -216 ± 22 | -798 ± 65 |
| | C10H6 | 71.3 ± 1.4 | 642 ± 21 | 1841 ± 64 | 65 ± 0.2 | 100.3 ± 1.3 | 382 ± 19 | 908 ± 56 | 29. ± 1.3 | -260 ± 20 | -933 ± 59 |
| | C12H6 | 77.6 ± 1.1 | 723 ± 17 | 2082 ± 50 | 65 ± 0.1 | 104.2 ± 1.3 | 434 ± 18 | 1064 ± 55 | 26.6 ± 1.3 | -289 ± 19 | -1018 ± 58 |
| $caC_{7.0}$ | C2H6 | 75.4 ± 7.0 | 694 ± 104 | 1993 ± 312 | 65.2, ± 1.0 | 98.7, ± 7.8 | 366 ± 115 | 862 ± 346 | 23.3, ± 8.2 | -328 ± 121 | -1132 ± 365 |
| | A4H2 | 72.3 ± 1.5 | 605 ± 23 | 1718 ± 68 | 67.7 ± 0.3 | 107.3 ± 1.8 | 462 ± 26 | 1145 ± 78 | 35.0 ± 1.7 | -143 ± 25 | -573 ± 74 |
| | T5H6 | 75.0 ± 0.9 | 696 ± 14 | 2002 ± 42 | 64.8 ± 0.1 | 106.6 ± 1.1 | 481 ± 16 | 1208 ± 49 | 31.6 ± 1.1 | -215 ± 16 | -794 ± 48 |
| | A8H2 | 78.5 ± 2.0 | 701 ± 29 | 2006, ± 87 | 66.4 ± 0.3 | 106.3 ± 1.6 | 445 ± 23 | 1092 ± 70 | 27.8 ± 1.9 | -256 ± 27 | -914 ± 81 |
| | A8H8 | 73.6 ± 1.4 | 663 ± 21 | 1899 ± 63 | 65.5, ± 0.2 | 107.9 ± 2. | 485 ± 29 | 1217 ± 88 | 34.3 ± 2.1 | -177 ± 30 | -682 ± 91 |
| | T9H6 | 75.7 ± 0.8 | 700 ± 13 | 2014 ± 38 | 65., ± 0.1 | 99.2 ± 0.9 | 372 ± 13 | 879 ± 40 | 23.4 ± 0.9 | -328 ± 13 | -1134 ± 40 |
| | C10H6 | 77.3 ± 3.6 | 727 ± 59 | 2096 ± 180 | 64.7, ± 0.6 | 98.6 ± 3.2 | 371 ± 51 | 879 ± 156 | 21.3 ± 2.9 | -356 ± 46 | -1217 ± 140 |
| | C12H6 | 81.8 ± 2.4 | 785 ± 38 | 2267 ± 114 | 64.6, ± 0.3 | 99.6 ± 1.7 | 374 ± 26 | 884 ± 80 | 17.7 ± 1.9 | -411 ± 30 | -1382 ± 91 |
| $caC_{5.8}$ | C2H6 | 66.5 ± 1.5 | 591 ± 23 | 1691 ± 68 | 64.9 ± 0.3 | 94.4 ± 1.8 | 318 ± 27 | 722 ± 81 | 27.9 ± 1.7 | -273 ± 26 | -969 ± 79 |
| | A4H2 | 64.4 ± 1.1 | 515 ± 18 | 1452 ± 53 | 67.9 ± 0.3 | 97.6 ± 1.2 | 333 ± 18 | 759 ± 54 | 33.2 ± 1.2 | -182 ± 19 | -693 ± 57 |
| | A4H8 | 73. ± 1.7 | 690 ± 27 | 1991 ± 80 | 64. ± 0.2 | 101.4 ± 2.3 | 418 ± 34 | 1022 ± 104 | 28.4 ± 2.1 | -272 ± 33 | -969 ± 98 |
| | T5H6 | 60.7 ± 2.3 | 519 ± 35 | 1478 ± 106 | 64.9 ± 0.4 | 85.5 ± 2.6 | 195 ± 40 | 354 ± 121 | 24.8 ± 2.1 | -324 ± 32 | -1125 ± 97 |
| | A8H2 | 71.6 ± 1.3 | 628 ± 19 | 1795 ± 58 | 66. ± 0.2 | 97.9 ± 1.2 | 343 ± 19 | 792 ± 56 | 26.4 ± 1.1 | -285 ± 18 | -1003 ± 53 |
| | A8H8 | 68.4 ± 1.2 | 615 ± 18 | 1761 ± 53 | 64.8 ± 0.2 | 102.7 ± 1.5 | 430 ± 22 | 1056 ± 67 | 34.2 ± 1.5 | -185 ± 23 | -705 ± 69 |
| | T9H6 | 65.2 ± 1.2 | 574 ± 18 | 1640 ± 54 | 64.9 ± 0.2 | 94.9 ± 1.1 | 328 ± 17 | 750 ± 53 | 29.7 ± 1.2 | -246 ± 18 | -889 ± 53 |
| | C10H6 | 67.4 ± 1.4 | 604 ± 22 | 1730 ± 66 | 64.8 ± 0.2 | 92.4 ± 1.2 | 290 ± 19 | 638 ± 56 | 24.9 ± 1.1 | -314 ± 17 | -1092 ± 53 |
| | C12H6 | 73.6 ± 1.1 | 682 ± 17 | 1961 ± 50 | 64.6 ± 0.2 | 95.8 ± 1.3 | 332 ± 19 | 762 ± 58 | 22.2 ± 1.1 | -350 ± 17 | -1199 ± 51 |
| $caC_{5.8}$ | C2H6 | 58.6 ± 0.7 | 483 ± 11 | 1369 ± 32 | 65.6 ± 0.2 | 93.1 ± 0.7 | 309 ± 11 | 695 ± 34 | 34.5 ± 0.9 | -175 ± 14 | -674 ± 42 |
| | A4H2 | 58.5 ± 0.5 | 441 ± 8 | 1232 ± 24 | 68.5 ± 0.2 | 96.5 ± 0.5 | 339 ± 7 | 782 ± 22 | 38. ± 0.4 | -102 ± 7 | -451 ± 21 |
| | A4H8 | 64.9 ± 0.7 | 583 ± 11 | 1669 ± 34 | 64.3 ± 0.1 | 96.6 ± 0.8 | 357 ± 12 | 840 ± 37 | 31.8 ± 0.8 | -226 ± 12 | -829 ± 35 |
| | T5H6 | 60. ± 0.4 | 519 ± 6 | 1479 ± 19 | 64.4 ± 0.1 | 94.2 ± 0.5 | 324 ± 8 | 740 ± 24 | 34.3 ± 0.6 | -195 ± 9 | -739 ± 26 |
| | A8H2 | 64.5 ± 0.6 | 546 ± 9 | 1552 ± 27 | 66. ± 0.1 | 98.1 ± 0.6 | 368 ± 9 | 870 ± 29 | 33.6 ± 0.6 | -178 ± 9 | -682 ± 27 |
| | A8H8 | 60.8 ± 0.5 | 505 ± 8 | 1433 ± 25 | 65.8 ± 0.1 | 92.3 ± 0.5 | 294 ± 8 | 650 ± 24 | 31.5 ± 0.5 | -211 ± 8 | -783 ± 23 |
| | T9H6 | 62.1 ± 0.4 | 550 ± 7 | 1575 ± 21 | 64.1 ± 0.1 | 95.2 ± 0.4 | 347 ± 7 | 813 ± 20 | 33.1 ± 0.5 | -203 ± 7 | -761 ± 21 |
| | C10H6 | 56.8 ± 0.6 | 454 ± 10 | 1280 ± 29 | 66.1 ± 0.2 | 92.9 ± 0.6 | 312 ± 9 | 708 ± 26 | 36.1 ± 0.6 | -142 ± 9 | -573 ± 28 |
| $fC_{7.0}$ | G1H8 | 64.6 ± 7.4 | 470 ± 128 | 1306 ± 389 | 75 ± 14.1 | 86.8 ± 4.6 | 122 ± 79 | 115 ± 241 | 22.2 ± 4.7 | -347 ± 80 | -1191 ± 244 |
| | C2H6 | 74.7 ± 10.4 | 769 ± 158 | 2238 ± 475 | 62 ± 0.5 | 99.7 ± 5.6 | 413 ± 84 | 1011 ± 253 | 25. ± 5.5 | -356 ± 83 | -1227 ± 250 |
| | A4H2 | 73.1 ± 1. | 712 ± 16 | 2060 ± 49 | 63 ± 0.1 | 101.6 ± 0.6 | 432 ± 9 | 1064 ± 28 | 28.5 ± 0.6 | -281 ± 9 | -996 ± 28 |
| | A4H8 | 71.9 ± 1.2 | 716 ± 19 | 2078 ± 58 | 62 ± 0.1 | 101.7 ± 1.2 | 444 ± 18 | 1102 ± 55 | 29.8 ± 1.1 | -273 ± 18 | -975 ± 53 |
| | T5H6 | 64.3 ± 1.1 | 613 ± 18 | 1768 ± 53 | 62 ± 0.1 | 93.9 ± 0.8 | 340 ± 12 | 792 ± 38 | 29.6 ± 0.8 | -273 ± 12 | -976 ± 37 |
| | C6H6 | 67.4 ± 1.3 | 652 ± 20 | 1886 ± 60 | 63 ± 0.1 | 99.9 ± 1.6 | 422 ± 24 | 1038 ± 73 | 32.5 ± 1.2 | -230 ± 19 | -848 ± 58 |
| | C6H7 | 48.6 ± 1.6 | 344 ± 26 | 953 ± 78 | 68 ± 0.8 | 118.8 ± 1.8 | 640 ± 28 | 1680 ± 86 | 70.2 ± 1.9 | 296 ± 30 | 727 ± 90 |
| | A8H2 | 71.2 ± 0.6 | 678 ± 9 | 1955 ± 28 | 64 ± 0.1 | 99.6 ± 0.4 | 398 ± 6 | 962 ± 18 | 28.4 ± 0.4 | -280 ± 6 | -994 ± 18 |
| | A8H8 | 65.3 ± 0.9 | 604 ± 14 | 1737 ± 42 | 63 ± 0.1 | 98.2 ± 0.7 | 383 ± 11 | 920 ± 34 | 32.9 ± 0.6 | -221 ± 10 | -817 ± 31 |
| | T9H6 | 64.6 ± 1.2 | 611 ± 18 | 1761 ± 55 | 63 ± 0.1 | 97.1 ± 1. | 383 ± 16 | 920 ± 47 | 32.5 ± 1. | -228 ± 15 | -841 ± 45 |
| | C10H6 | 66.4 ± 1.2 | 636 ± 19 | 1835 ± 57 | 63 ± 0.2 | 97.5 ± 1. | 384 ± 16 | 924 ± 49 | 31.1 ± 1.1 | -252 ± 17 | -911 ± 50 |
| | C12H6 | 69.7 ± 1. | 679 ± 15 | 1964 ± 47 | 63 ± 0.1 | 98.7 ± 0.9 | 402 ± 14 | 977 ± 43 | 29. ± 1. | -277 ± 15 | -987 ± 45 |
